# A protective bispecific antibody targets both Nipah virus surface glycoproteins and limits viral escape

**DOI:** 10.1101/2025.03.11.642517

**Authors:** Ariel Isaacs, Guillermo Valenzuela Nieto, Xinghai Zhang, Naphak Modhiran, Jennifer Barr, Nazia Thakur, Yu Shang Low, Rhys H. Parry, James B. Barnes, Ronald Jara, Johanna Himelreichs, Yanfeng Yao, Camila Deride, Barbara Barthou-Gatica, Constanza Salinas-Rebolledo, Ehrenfeld Pamela, Jun Jet Hen, Noah Hayes, Devina Paramitha, Mahali S. Morgan, Christopher L.D. McMillan, Martina L. Jones, Trent Munro, Alexander A. Khromykh, Patrick C. Reading, Paul R. Young, Keith Chappell, Yi Shi, Dalan Bailey, Glenn Marsh, Sandra Chiu, Alejandro Rojas-Fernandez, Daniel Watterson

**Author notes:** These authors contributed equally (Ariel Isaacs, Guillermo Valenzuela Nieto, Xinghai Zhang).

## Abstract

Nipah virus (NiV) and Hendra virus (HeV) are highly pathogenic henipaviruses without approved human vaccines or therapies. Here, we report on a highly potent bispecific therapeutic that combines an anti-fusion (F) nanobody with an anti-receptor binding protein (RBP) antibody to deliver a dual-targeting biologic that is resistant to viral escape. We show that the nanobody, DS90, engages a unique, conserved site within prefusion F of NiV and HeV, and provides neutralization and complete protection from NiV disease. Bispecific engineering of DS90 with the anti-RBP mAb m102.4 results in neutralization, elimination of viral escape and superior protection from NiV disease compared to leading monovalent approaches. These findings carry implications for the development of cross-neutralizing immunotherapies that limit the emergence of henipaviral escape mutants.

## Main

Nipah virus (NiV) and Hendra virus (HeV) are the prototypical members of the *Henipavirus* (HNV) genus and are regarded as the most pathogenic paramyxoviruses^1,2^. Infection with NiV and HeV presents as severe respiratory symptoms and progresses on to febrile encephalitis, with an associated mortality rate of 50-100%^3^. The *Pteropus* fruit bats are the main animal reservoir of HNVs, yet infection can occur in animals from at least six mammalian orders. Since their emergence in the 1990’s, numerous spillover events of HeV from bats to horses have occurred in Australia, resulting in several human exposures^4^. NiV spillovers to humans occur almost annually in Bangladesh, and several outbreaks have also been recorded in India and the Philippines, consistent with the range of the reservoir host^3^.

There are currently no approved human vaccines or therapeutics for HNVs. Until recently, the primary therapeutic and vaccine target has been the receptor binding protein (RBP, previously designated G), a type II homotetrameric glycoprotein that mediates attachment to host cells through engagement with ephrin B2/B3 receptors^5,6^. This has led to the development of the leading biologic, m102.4 - a monoclonal antibody (mAb) that targets the receptor binding site within RBP and neutralizes both NiV and HeV^7^. This antibody has shown success in several animal challenge models and was found to be well tolerated in a human phase I clinical trial^8–12^.

An alternate therapeutic target is the HNV fusion (F) glycoprotein. HNV F is a class I homotrimeric protein that is initially synthesized as an F_0_ precursor^13,14^. Similarly to other paramyxovirus fusion glycoproteins, HNV F initially exists as a poised, metastable prefusion trimer ^13,14^. After attachment and triggering by RBP, F undergoes a series of conformational changes from its prefusion form to a highly stable post-fusion form, driving the fusion of viral and host membranes. Animal immunizations with prefusion F have led to the isolation of several potent neutralizing antibodies^15–18^. A humanized version of a prefusion F-specific mAb 5B3 was recently found to be protective in animal challenge models, providing a basis for the development of F-based therapeutics^19^.

Despite this work, antibody therapy against viral infections meets several limitations. RNA viruses, such as HNVs, frequently mutate under antibody selection pressures, leading to the emergence of resistance. This has been most evident in therapies targeting SARS-CoV-2 spike, where several antibody therapeutics lost their efficacy against newly emerged viral variants^20–22^. For HNVs, several studies have demonstrated that single amino acid substitutions in the RBP or F completely ablate the neutralization of leading mAbs such as m102.4 and 5B3^16,23,24^. To address this, bispecific antibodies that target RBP have been trialed, yet have not demonstrated superior protective efficacy over monovalent treatments^25^. Furthermore, antibody cocktails targeting distinct epitopes within F have been shown to be less protective than a single antibody treatment^26^. In comparison, RBP cocktails have provided synergism^25^, but given the higher genetic diversity observed for RBP compared to F, an ideal therapeutic would target neutralizing sites on both surface antigens and offer protection from disease and resistance to viral escape^27,28^. Here, we address this directly by engineering the first dual-targeting bispecific antibody that recognizes neutralizing sites within both F and RBP. To achieve this, we first set out to discover novel F-targeting VHH therapeutics, which when fused to the clinically tested anti-RBP m102.4 could offer an elegant solution, enabling dual-antigen targeting together with bivalent presentation and antibody effector functionality.

## Results

### Nanobodies targeting NiV & HeV F offer ultrapotent neutralizing activity

NiV-Malaysia (NiV-M) F-specific nanobody sequences were isolated from alpaca peripheral blood mononuclear cells (PBMCs) immunised with prefusion stabilized NiV-M F using an approach previously established for SARS-CoV-2 nanobody discovery^29^ (Figure 1A). VHH genes were amplified to generate a nanobody bacterial display library with a size of 6.2 × 10^6^ individual clones with <1% of re-ligated vectors. This library was then incubated with NiV-M F-coated Sepharose beads to pull down bacteria expressing F-specific nanobodies via a Ficoll-based density gradient, where we obtained ∼150 colonies on LB-agar plates from the Sepharose-antigen coated fraction (Figure S1A). Bacterial lysates from isolated clones were initially screened by immunofluorescence assay against NiV F expressing cells to confirm binding and specificity (Figure 1A, Figure S1B). Four clones, DS29, DS34, DS76 & DS90, co-localised with NiV-M F expressing cells (under pIRES-GFP) and were subsequently sequenced (Figure S1C). This panel of nanobodies was recombinantly expressed as dimers (fused to human Fc) or monomers (fused to human monomeric Fc (FcM)) in the ExpiCHO system. All nanobodies expressed well in both formats, with the exception of DS29, where low yield limited downstream analyses. Affinity purification resulted in highly pure nanobody Fc dimers and FcM, as visualised by non-reducing SDS-PAGE (Figure S1D,E). Next, we confirmed binding by both surface plasmon resonance (SPR) and ELISAs of nanobody monomers and dimers against both NiV-M F and HeV F glycoproteins. We obtained SPR dissociation constants (K_D_) against NiV F of 4.83 nM (DS90), 23.4 nM, (DS34), 303 nM (DS76) when expressed as monomers (Figure 1B). In an ELISA, nanobody Fc dimers bound to prefusion HeV F with apparent affinities of 1 nM (DS90), 11 nM (DS34) and 14 nM (DS76) (Figure 1B, Figure S1D,E). Due to low expression levels, the affinity of DS29 Fc dimer was not tested. The monomeric DS90 was also expressed recombinantly by periplasmic expression in *E. coli*, tested for immunofluorescence and was shown to be thermally stable by ELISA (Figure S1F-H).

**Figure 1.**
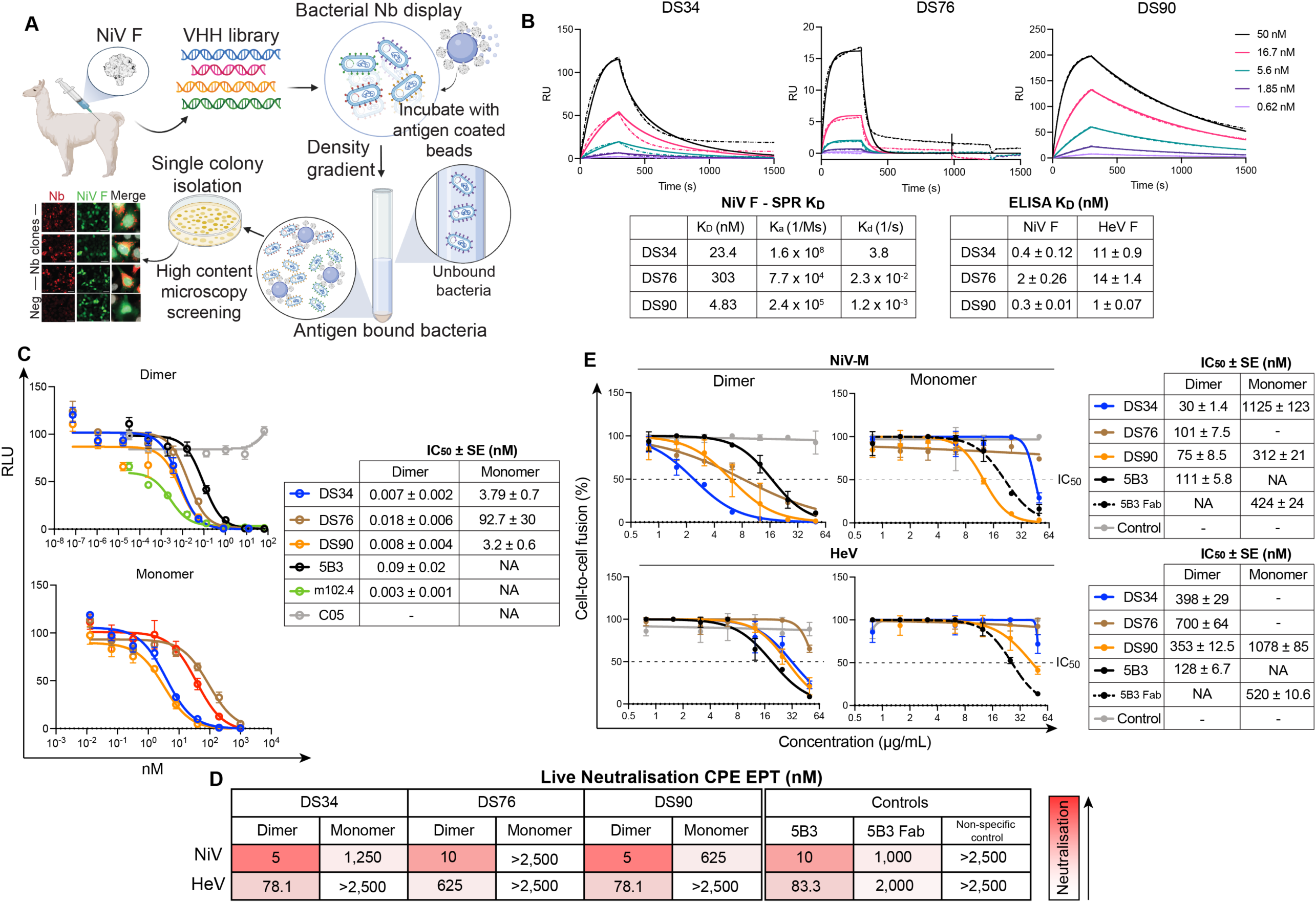
Isolation and functional characterisation of F-specific nanobodies. (A) Summarised workflow for nanobody discovery. An alpaca was immunised with prefusion NiV F prior to generation of VHH bacterial display library. NiV F-specific nanobodies are enriched using a density gradient and screened via high-resolution confocal microscopy (Figure S1B). (B) SPR sensorgrams showing binding between NiV F and DS34, DS76 and DS90 Fc monomers. Binding and fitted curves are shown as dotted and solid lines, respectively. SPR derived dissociation constants (K_D_) are shown below. ELISA derived K_D_ are shown against prefusion NiV and HeV F proteins (binding curves are shown in Fig S1D-E). (C) NiV pseudovirus neutralization curves of nanobody dimer (Fc) or monomers (FcM) benchmarked against known neutralizing mAbs and a non-specific isotype control (C05). Data shown is an average of triplicates with standard deviation and is normalised to virus only control. IC_50_s with standard error are shown to the right. (D) Authentic NiV neutralization by nanobody Fc dimer or monomers conducted in BSL4 shown as cytopathic effect endpoint titre (CPE EPT). Data shown is an average of duplicate values. (E) Micro-fusion inhibition tests (mFIT) of nanobody Fc dimers or monomers against NiV (top) and HeV (bottom). Data shown is an average of triplicates with standard deviation. Dashed lines represent 50% inhibition and are shown as IC_50_ values with standard error to the right.

To assess functionality, we conducted neutralization assays in a NiV pseudotyped assay, where both F and RBP are presented on lentiviral virions (Figure 1C). Here, we found that all nanobody constructs were able to neutralize NiV pseudovirus to various extents, with DS34 and DS90 Fc dimers achieving the highest neutralization with an IC_50_ of 7-8 pM. Further comparisons of DS90 Fc dimer neutralisation capacity were made in the pseudovirus assay with a larger panel of previously described antibodies against NiV-M and NiV Bangladesh-2008 strain (NiV-B08) (Figure S2a-c) ^15–18^. Here, DS90 Fc dimer was highly neutralising against both NiV-M and NiV-B08 with IC_50_s of 0.007 and 0.002 nM, respectively. DS90 Fc dimer was the most potent neutraliser for both strains across the whole panel, and only 4H3 achieved similar levels against NiV-M (IC_50_ = 0.009 nM), but had a reduced neutralisation against NiV-B08 by 8.5-fold compared with DS90 Fc dimer. To confirm the pseudovirus neutralisation results, we also conducted a neutralization assay in Vero cells against authentic NiV-M & HeV under BSL4 conditions (Figure 1D). We found that both DS34 and DS90 Fc dimers were able to neutralize both NiV-M and HeV with cytopathic effect endpoints titres (CPE EPT) of 5 nM and 78 nM respectively. We also tested DS90 Fc dimer neutralisation against the NiV-B-2004 strain (NiV-B04) in a plaque reduction neutralisation test where it had an IC_50_ of 2.4 nM, surpassing that of 5B3 comparator which had limited neutralisation ability against NiV-B04 (Figure S2D).

To further investigate the mechanism underpinning the neutralization offered by the nanobodies, we conducted micro-fusion inhibition tests (mFITs) to determine fusion inhibition efficacy against both NiV-M and HeV (Figure 1E). We found that DS90 & DS34 Fc dimers were potent inhibitors of NiV cell fusion with IC_50_s of 75 nM and 30 nM, respectively. Despite only one amino acid difference in complementarity determining region (CDR) 3 between DS90 and DS34 (V101A), DS34 Fc was able to inhibit fusion to a higher degree than DS90 Fc, however this was significantly reduced as a monomer (Figure 1E). In line with the SPR and ELISA binding data, all constructs were less effective at inhibiting HeV cell fusion.

### DS90 binds the lateral face of NiV F and engages a novel quaternary pocket

Based on functional data, we progressed DS90 as the lead nanobody candidate for epitope elucidation via cryogenic electron microscopy (cryoEM) (Figure 2). Soluble DS90 VHH domains were produced through proteolytic cleavage of a HRV3C site at the Fc region, and purified by size-exclusion chromatography prior to complexation with NiV F (Figure S3A,B). Separated DS90-complexed NiV F was concentrated and plunge frozen for cryoEM imaging. Using single particle analysis (SPA), we determined a cryoEM structure of NiV F complexed with DS90 to 3.66 Å resolution (FSC 0.143, Figure 2, Figure S4, Table S1).

**Figure 2.**
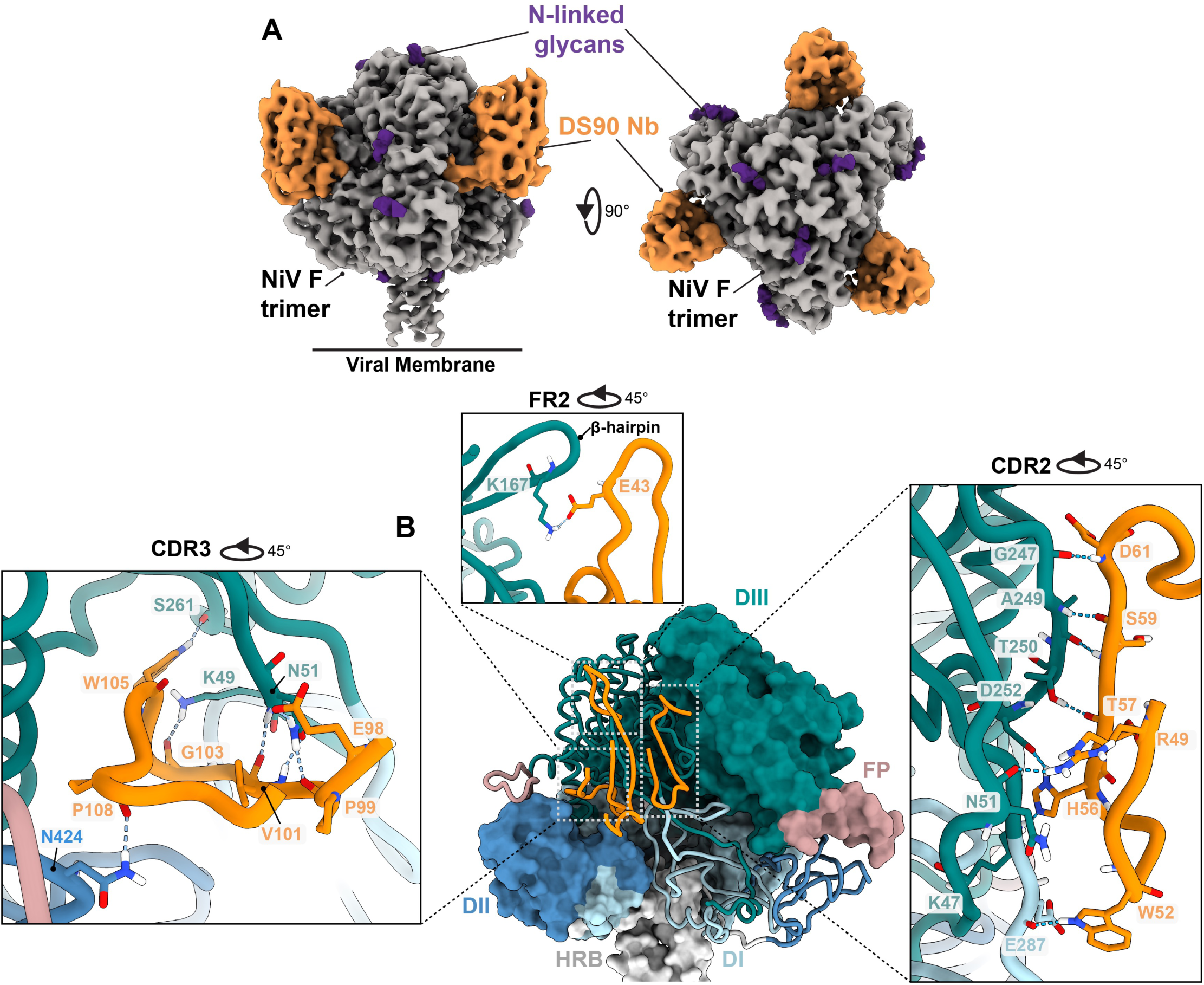
DS90 binds to a novel, glycan-free, quaternary epitope on NiV. **F.** (A) CryoEM map of three DS90 nanobodies (orange) bound to NiV prefusion F trimer (grey) with N-linked glycans shown in purple. (B) Structural representation of NiV F trimer with domains coloured light cyan (DI), blue (DII), teal (DIII) and coral (fusion peptide). DS90 loops that contribute to the binding interface are shown (orange licorice representation). Insets show a rotated view of hydrogen bonds and salt bridge contacts CDR2, CDR3 and framework region 2 (FR2) make with NiV F trimer. Side chains of residues that form contacts are shown in stick representation, with CPK heteroatom colouring.

DS90 binds to a glycan-free site on the lateral face of NiV F trimer (Figure 2A). The DS90 epitope is quaternary and spans DI and DIII within one NiV F protomer and contact DII of the adjacent F promoter (Figure S3C). Binding is coordinated predominantly by CDRs 2 & 3, which together constitute 1428.3 Å^2^ buried surface area spanning two NiV F protomers (Figure 2B). CDR2 of DS90 contacts the connecting strands of NiV F that link DI and DIII, mediated by hydrogen bonds between H56_DS90_ and K47_F_,W52_DS90_ and E287_F_, as well as the CDR2 adjacent FR residue R49_DS90_ which forms a hydrogen bond with N51_F_ (Figure 2B, Figure S5A-B, Table S2). CDR2 also interacts with a central loop region of DIII on the lateral face of the F trimer through hydrogen bonds formed between H56_DS90_ and D252_F_, T57_DS90_ and T250_F_, S59_DS90_ and A249_F_ and D61_DS90_ and G247_F_.

The long CDR3 of DS90 projects into the cavity formed by DI, DIII and the fusion peptide of one protomer and DII of the adjacent protomer (Figure 2B, Figure S5C-D). Recognition is in-part mediated by E98_DS90_, P99_DS90_ and V101_DS90_, which all make hydrogen bonds with N51_F_ on the outer face of the pocket formed between two neighbouring F protomers (Figure S3C, Table S2). The looping CDR3 wraps into the interprotomer pocket, stabilized by a hydrogen bond between the carbonyl group of G103_DS90_ and K49_F_. Deep within the cavity, the bulky side chains of W105_DS90_ and F106_DS90_ sit within the hydrophobic patch formed by the partially hidden fusion loop. W105_DS90_ also forms a hydrogen bond with S261_F_. Notably, the backbone carbonyl of P108_DS90_ forms a hydrogen bond with N424_F_ of a neighbouring F protomer, thus stabilizing the prefusion F trimer at the quaternary level (Figure 2B). N424 sits proximal to a β-strand succession previously shown to be important for sequestering the fusion peptide and securing the prefusion state of F (Figure S6) ^30^.

Interestingly, comparisons with the apo-form of NiV F (PDB 5EVM) and other mAb-bound NiV F structures (PDB 6TYS & 7UPD) suggested that the interaction with P108_DS90_ causes a shift in the position and orientation of N424_F_ side chain (Figure S6A). To confirm this, we performed symmetry expansion and a local refinement on this region (Figure S5B). Here, we observed the map density of N424 side chain to be shifted facing outward compared with a previously determined cryoEM structure of apo-F (PDB 8DNG), which faces inwards towards the trimer (Figure S6C). As such, this altered conformation, mediated by DS90, may provide further stabilisation of the fusion peptide region within a single F trimer.

Further structural investigation revealed that the region of NiV F that forms contacts with CDR3 is involved in the previously reported F dimer-of-trimer assembly, where the hydrophobic fusion peptide of one F trimer is sequestered in the dimer-of-trimer interface (Figure S6D). The biological function of the higher oligomeric assembly of NiV F remains to be explored, yet is thought to be crucial for coordination of viral membrane fusion^30,31^. As such, CDR3 of DS90 may provide a functional mimic of the adjacent fusion peptide, thereby disrupting the higher-order oligomer prefusion F assembly and preventing coordinated viral membrane fusion events.

In addition to the CDRs, DS90 also forms contacts with NiV F through residues in FR2, however, lower local resolution at this region prevents a definitive description of this interaction (Figure S5E). Here, FR2 likely contacts a β-hairpin within the NiV F heptad repeat A (HRA) via a salt bridge between E43_DS90_ and K167_F_ (Figure 2B, Figure S3C, Table S2). This is tentatively strengthened by additional Van der Waals interactions between G41_DS90_ & K42_DS90_ with E166_F_ of the HRA (Table S3).

Structural comparisons with previously characterized murine-derived mAbs revealed that DS90 binds a distinct site relative to apical (12B2, mAb66, 4H3) and basal (1H1, 2B12) targeting mAbs (Figure S7), although there is some overlap. Notably, the epitope of apical mAb 2D3 includes the β-hairpin of the NiV F HRA which also makes contact with DS90 FR2. Epitope comparisons also show significant overlap with mAbs that bind the lateral face of NiV F including 1H8, 1A9, 5B3 and 1F5 (Figure S7) ^15–18^. Despite this, the elongated CDR3 of DS90 forms a unique interaction within the deep quaternary pocket adjacent to the fusion peptide that has not been seen in previous mAb-F interactions.

The DS90 epitope is largely conserved between NiV-M and HeV F proteins, with only a single semi-conserved substitution at T263A (Figure S3C, Figure S8A). Furthermore, two NiV-B strains contain substitution mutations at either D252G (NiV-B/2004) or A249T (NiV-B/2012), which form hydrogen bonds with DS90 (Figure S8A). We previously demonstrated that DS90 Fc dimer is able to neutralise NiV-B04 (containing D252G) (Figure S2D). To investigate the effect of A249T and T263A mutations, we produced pseudoviruses containing these mutations within NiV-M F backbone and conducted neutralisation tests (Figure S8B). Our results demonstrate minimal change in IC_50_ between wild-type NiV pseudovirus and NiV F-A249T or F-T263A mutants, suggesting that these substitution events do not confer resistance to DS90 neutralisation.

### A bispecific therapeutic strategy targets both F and RBP glycoproteins

Nanobodies present as a promising strategy to combat HNV infection and disease. However, HNVs are RNA viruses and therefore exhibit a high mutation rate, which may lead to immune evasion. Indeed, we and others have demonstrated that two amino acid substitutions in F compromise the neutralization offered by 5B3 or mAb66^16,32^. It has also been previously demonstrated that a single amino acid substitution in HNV RBP results in decreased neutralization by leading therapeutic mAb m102.4^23,24^. To address this, we sought to generate a bispecific therapeutic strategy that would target both F and RBP and therefore limit the emergence of viral escape mutants. Additionally, bispecific antibodies may provide increased or synergistic virus neutralization effects over monoclonal antibodies, and their design as single-molecule biologics streamlines their production and reduces costs in comparison to antibody cocktails^33–35^. To leverage these advantages, we engineered a bispecific antibody sconsisting of DS90 genetically fused to the N-terminus of m102.4 mAb heavy chain (DS90-m102.4) and confirmed its purity and homogeneity by SDS-PAGE and size-exclusion chromatography (Figure 3A, Figure S9A,B). Next, we evaluated binding to NiV F and RBP by both ELISA and SPR, and observed a similar binding capacity of DS90-m102.4 as previously measured for DS90 Fc and m102.4 (Figure 3B, Figure S9C). To determine whether DS90-m102.4 is able to simultaneously engage with F and RBP, we conducted a tandem SPR experiment where cumulative binding can be measured during independent F and RBP association phases (Figure 3C). Here, our results demonstrated that DS90-m102.4 is able to bind both NiV F and RBP simultaneously, allowing for cooperative binding.

**Figure 3.**
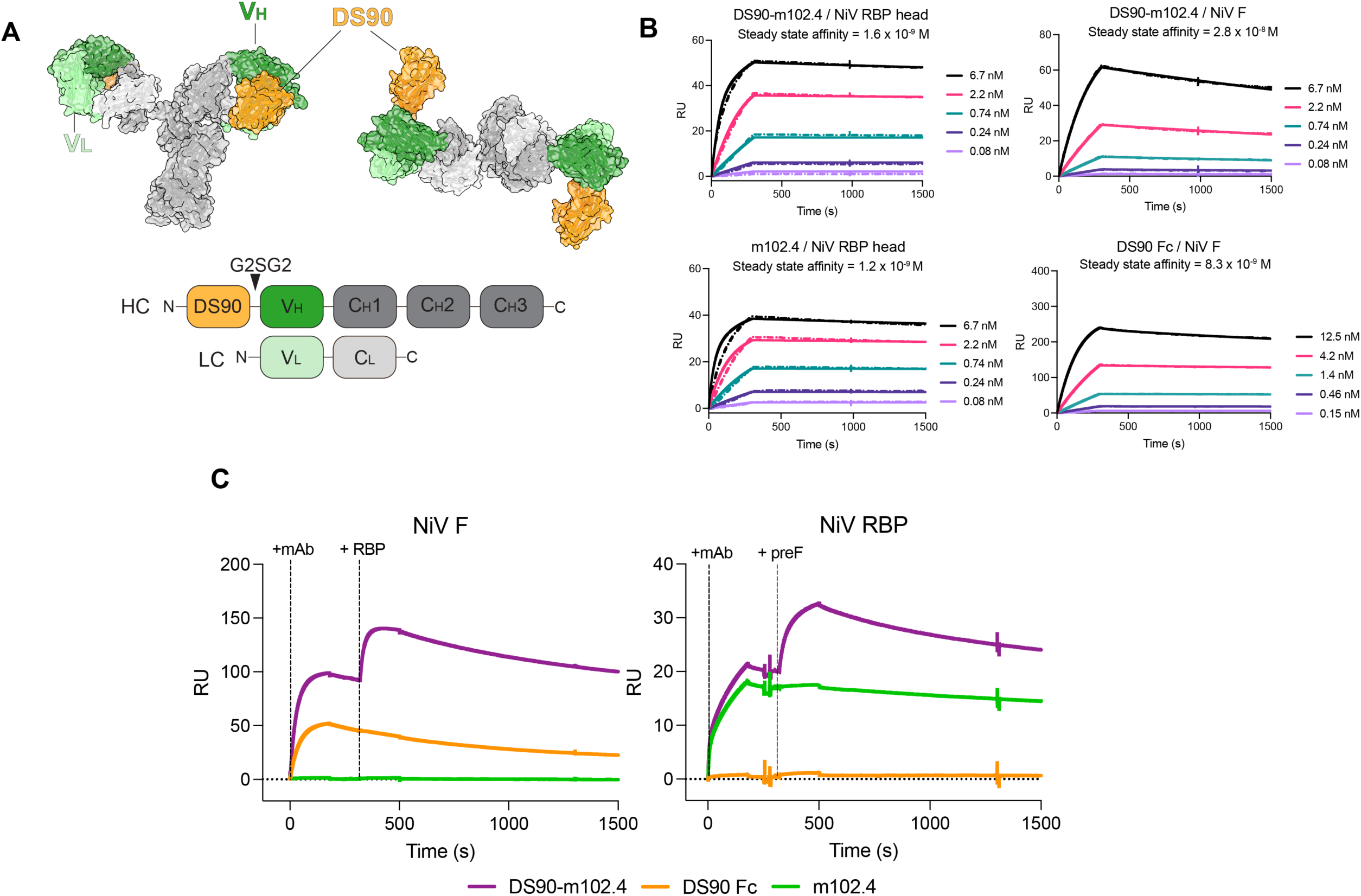
Design and characterisation of a bispecific DS90-m102.4 therapeutic approach. (A) Hypothetical 3D structure of the DS90-m102.4 bispecific antibody, modelled on m102.4 (PDB 6CMI). DS90 is coloured in orange and m102.4 variable heavy (V_H_) and variable light (V_L_) are coloured dark and light green, respectively. m102.4 constant domains are coloured grey. The DS90 VHH domain is added on to the N-terminus of the m102.4 heavy chain, linked by a G_2_SG_2_ linker. (B) SPR sensorgrams showing DS90-m102.4 binding to both NiV F and RBP. Binding and fitted curves are shown as dotted and solid lines, respectively. SPR steady state affinities are shown in nM, calculated from bivalent analyte general kinetics model. (C) Raw SPR sensorgrams showing cumulative binding of NiV F and RBP head domain by DS90-m102.4. Biotinylated NiV F or RBP head is coated on the SA-chip before an initial association phase with antibody (+mAb) followed by a second association phase with unbiotinylated antigen (+preF or RBP) and a long dissociation phase.

Next, we tested the neutralization capacity of DS90-m102.4 in live CPE endpoint neutralization assays. DS90-m102.4 displayed a marginal increase in neutralization compared to DS90 Fc (∼2-fold) and higher neutralization compared to m102.4 against both NiV-M and HeV (∼38-fold) (Figure S9D). To confirm this finding, we conducted plaque neutralization assays in an independent laboratory against NiV-M, NiV-B04, and HeV (Figure 4A). DS90-m102.4 neutralized NiV-M, NiV-B04, and HeV with IC_50_ values of 2.2, 0.12 and 0.9 nM, respectively. This corresponds to a 2- and 4.5-fold improvement in neutralization against NiV-M and HeV compared with m102.4 (NiV-M & HeV IC_50_: 4.1 nM). Similar neutralisation of NiV-B04 was seen for DS90-m102.4 and m102.4 mAb (both with IC_50_ of 0.12 nM). Compared with DS90 Fc, DS90-m102.4 had a ∼5-fold increase in neutralisation of NiV-M (DS90 Fc IC_50_ of 10.6 nM; DS90-m102.4 IC_50_ of 2.2 nM) and similar neutralisation of HeV (DS90 Fc IC_50_ of 0.6 nM; DS90-m102.4 IC_50_ of 0.9 nM). Furthermore, DS90-m102.4 demonstrated a 20-fold improvement in neutralisation against NiV-B04 compared with DS90 Fc (DS90 Fc IC_50_ of 2.44 nM; DS90-m102.4 IC_50_ of 0.12 nM). Together, these results demonstrate an overall improved neutralisation profile for the bispecific DS90-m102.4 compared to its monospecific components against NiV-M, NiV-B04 and HeV.

**Figure 4.**
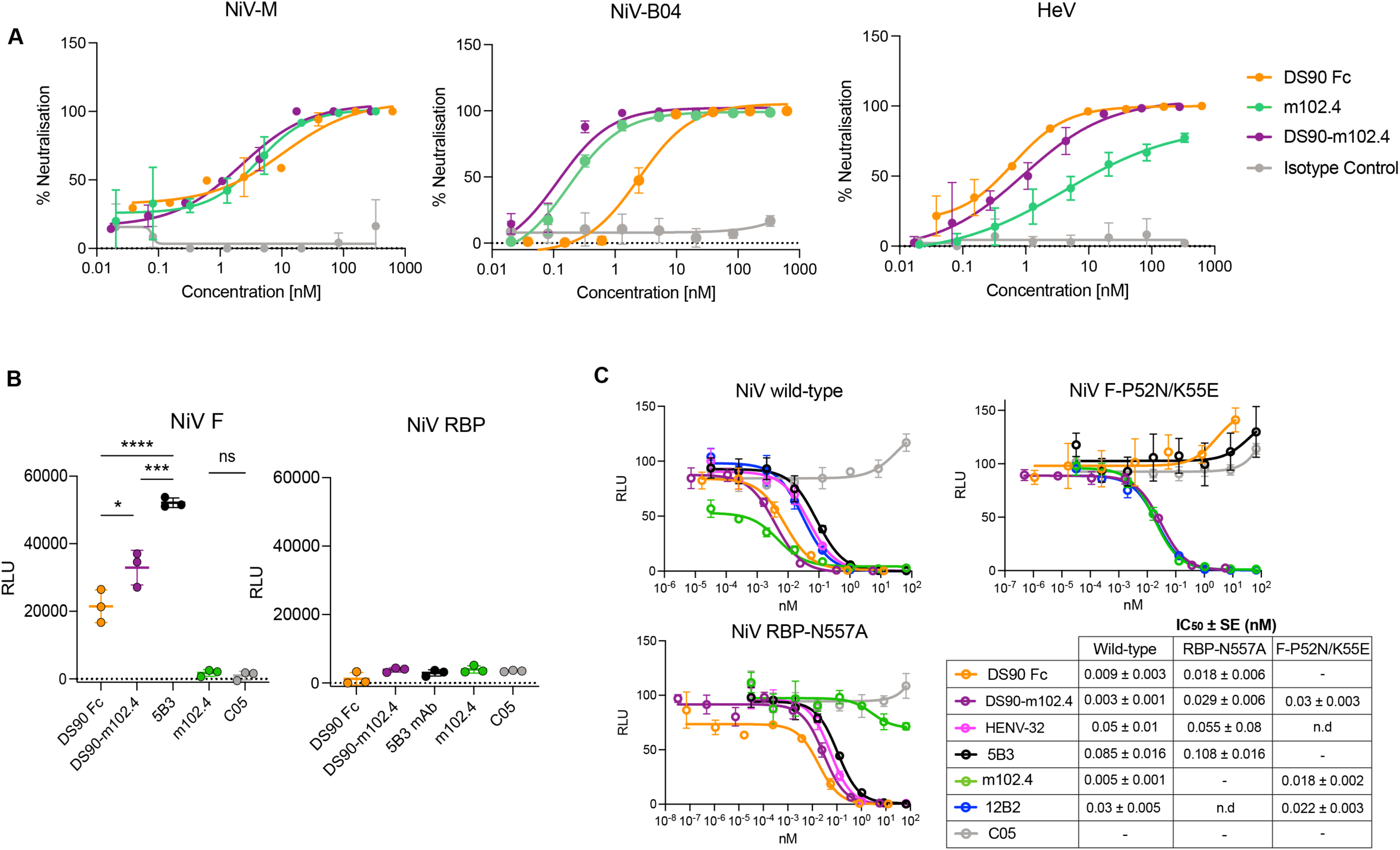
DS90-m102.4 provides effective neutralization against prospective escape mutants and offers ADCC effector functions. (A) Plaque reduction neutralisation tests of authentic NiV-M, NiV-Bangladesh/04 (NiV-B04) and HeV conducted at BSL4. Data shown is mean of triplicate values with standard deviation. Data was fit with a four parameter inhibitor vs. response model to quantify IC_50_ values. Non-specific antibody (C05) used as a negative isotype control. (B) ADCC effector functions of antibodies targeting NiV F and RBP measured at 20 μg/mL (associated data shown in Figure S10A). Data shown is mean of triplicate values with standard deviation. Statistical analysis conducted using an ordinary one-way ANOVA where * = p < 0.05, *** = p < 0.0005 and **** = p < 0.0001. (C) NiV neutralization tests of antibody constructs against pseudoviruses harbouring known F (P52N/K55E) and RBP (N557A) escape mutations. Data shown is of triplicate values with standard deviation. RLU data is normalised to virus only wells. IC_50_s are shown with standard error. Dashes indicate no neutralization observed and untested antibodies are denoted as not determined (n.d.).

While neutralization is a key factor in antiviral activity, protection from infection can also be mediated by other functions such antibody dependent cellular cytotoxicity (ADCC). To investigate this, stable HEK293T cells expressing either NiV F (293T-F) or RBP (293T-RBP) glycoproteins were established. The 293T-F or 293T-RBP were used as target cells in an ADCC reporter assay, mediated by effector Jurkat cells expressing NFAT-Luc upon activation. Here, we assessed the capacity of DS90 Fc, 5B3, m102.4 and DS90-m102.4 to elicit ADCC responses (Figure 4B, Figure S10A). Interestingly, it was observed that m102.4 did not induce ADCC effector functions despite binding to the 293T-RBP cells (Figure S10B). Conversely, 5B3 and DS90 Fc were able to activate ADCC function, with 5B3 providing significantly higher levels of induction. Furthermore, through addition of DS90 domains, the DS90-m102.4 bispecific antibody was able to induce F-specific ADCC functions. The bispecific approach therefore not only improves on live virus neutralization by m102.4, but also provides an ability to induce ADCC and therefore may offer other antiviral effects that provide protection from disease.

### DS90-m102.4 limits generation of NiV escape mutants

To further test the DS90-m102.4 bispecific antibody, we constructed NiV pseudovirus particles with mutations in either RBP or F (Figure 4C). Previous research has demonstrated that N557A mutation in NiV RBP ablated m102.4 binding^36^. Here, we confirmed that NiV pseudoviruses with this mutation could not be neutralized by m102.4; however, were permissible to neutralization by HENV-32, which binds an allosteric site on RBP to m102.4^37^. Here, the bispecific DS90-m102.4 antibody retained neutralization against this mutant with an IC_50_ of 0.03 nM, conferred by the DS90 domains.

Conversely, we tested the neutralization of several antibodies against pseudoviruses harbouring a previously described 5B3-resistant mutations in F (P52N/K55E)^16,32^ (Figure 4C). We first confirmed that 5B3 was unable to neutralize this mutant and also found that it was permissible to 12B2 mAb neutralization, which targets an alternate epitope^17^. NiV F-P52N/K55E pseudoviruses were also able to escape neutralization by DS90 Fc, yet were effectively neutralized by DS90-m102.4, with an IC_50_ of 0.03 nM. As such, by including both F and RBP targeting antibody domains within a single design, we were able to show efficient neutralization of prospective F and RBP escape mutants and therefore deliver a more resilient therapeutic approach.

We next aimed to demonstrate that DS90-m102.4 limits the likelihood of viral escape using live virus infection. Using authentic virus under BSL4 conditions, NiV was sequentially passaged in sub-neutralizing concentrations of antibody over 3 passages (P1-3) (Figure 5A). Concentrations of each antibody treatment was optimized for the selection conditions, where sub-neutralizing levels were defined as 2 µg/ml, 1 µg/ml and 0.2 µg/ml for m102.4, DS90 Fc and DS90-m102.4, respectively (Fig S11A&B). In the third passage (P3) the levels of DS90 Fc and DS90-m102.4 were increased to 2 and 0.5 µg/ml, respectively, to increase selection pressure. As anticipated, subsequent neutralization assays demonstrated NiV escape from neutralization by m102.4 and DS90 Fc (Figure S11C). However, virus passaged with DS90-m102.4 was effectively neutralized by m102.4, DS90 Fc and DS90-m102.4. To investigate the evolutionary determinants of viral escape in F and RBP genes under viral escape in the presence of treatment combinations, RT-PCR amplicons were generated from P3 viral RNA and then subjected to nanopore sequencing. The number of reads per sample ranged from 16,417 to 112,581. Reads were mapped to the Nipah virus NV/MY/99 isolate (GenbankID: AJ627196), and only reads that were greater than 500 nucleotides in length and with a mapping quality greater than 20 were retained. The resultant alignment files had an average per position coverage ranging from 3,548 to 23,528 allowing for examination of subconsensus viral populations. Following three passages in sub-neutralizing concentrations of m102.4 (2 μg/mL P1-3), sequencing data revealed that a single amino acid change within the RBP can ablate neutralization (Figure 5B, Figure S11D). This is in agreement with *in silico* designed single amino acid substitutions that knockout the m102.4 epitope^36^, further highlighting the ease by which NiV can escape neutralization. In our experiments, virus escape from m102.4 predominantly consisted of a S239L mutation within the RBP, found in 89% and 21% of reads of each replicate tested (Figure 5B, Figure S11D). Structural analysis revealed S239 to be positioned within the m102.4 epitope, however away from the receptor binding site, and makes key contacts with both the CDR3 and CDR2 of the m102.4 HC (Figure 5C). As such, mutation of this residue to a hydrophobic leucine likely ablates binding and mediates viral escape. An additional V587D mutation was observed within 78% of reads of a single replicate passaged in m102.4 (Figure S11D). While V587 sits in proximity to the m102.4 epitope, it does not make any direct contacts with the HC or LC. However, V587 resides within the B6 S2-3 loop of RBP, where changes have been previously noted to block m102.4 binding through allosteric mechanisms^24^. To validate these mutations, we generated NiV pseudoviruses with the RBP-S239L and RBP-V587D mutations. In pseudovirus neutralization assays, we observed a 23 and 57-fold reduction in neutralization capacity by m102.4 against the RBP-S239L and V587D mutants, respectively (Figure 5D). Interestingly, both of these escape mutations were recently corroborated by direct mutational scanning of RBP under m102.4 selection, and were found to not affect RBP receptor binding and viral entry^38^.

**Figure 5.**
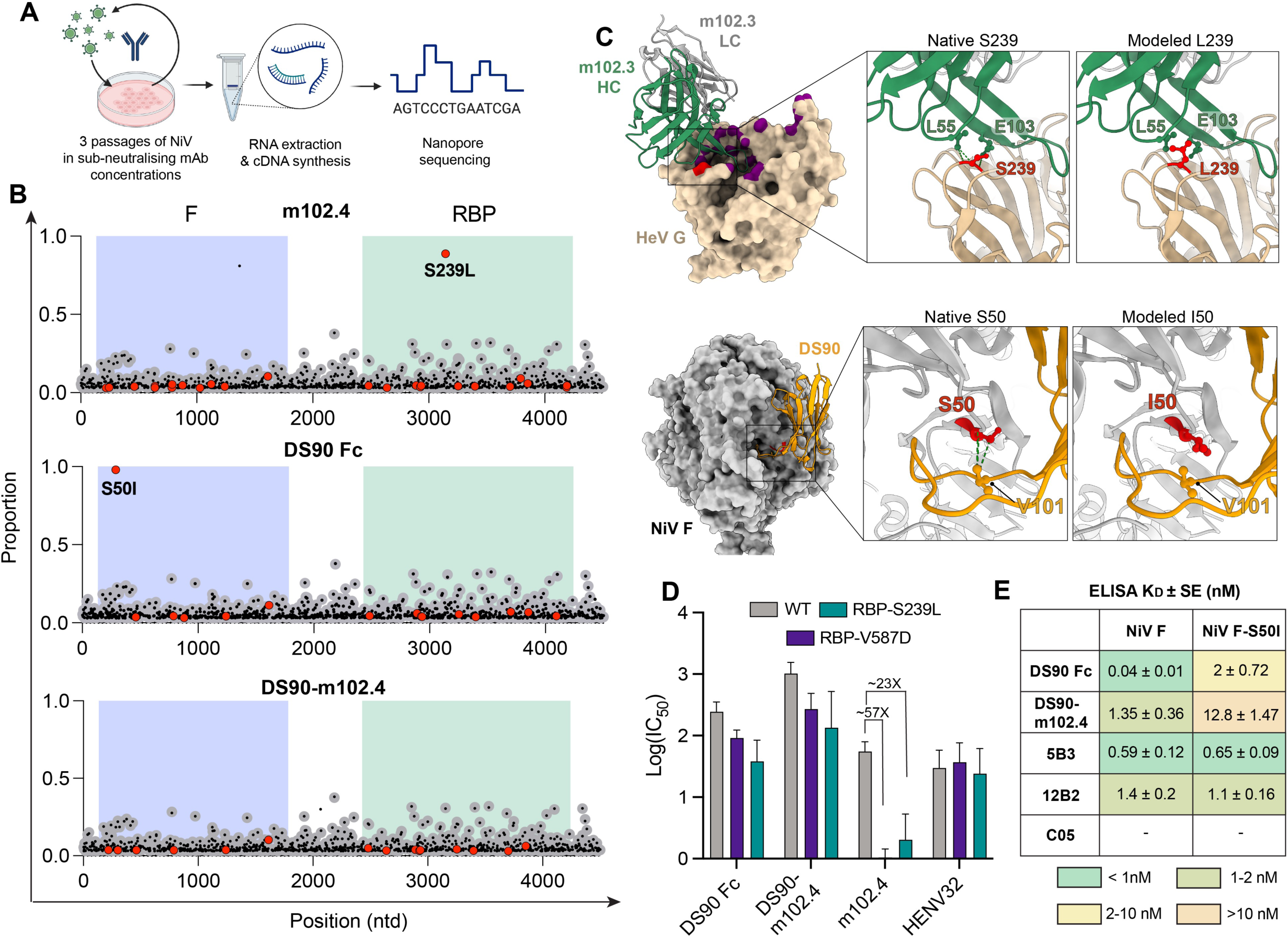
DS90-m102.4 limits the emergence of NiV escape mutants. (A) Experimental overview of NiV escape mutant generation. Authentic NiV is passaged thrice in sub-neutralizing concentrations of antibody. Total RNA is extracted and NiV F-RBP cDNA amplicon is subsequently generated for nanopore sequencing. (B) Incidence of escape mutations generated by antibody selection pressures within the F-RBP cDNA amplicon. Black dots represent synonymous mutations, grey circles represent stochastic mutations generated in the presence of a non-specific antibody (C05), red circles represent non-synonymous mutations. (C) Structural representation of S239L RBP mutation, shown using HeV RBP head structure bound to m102.3 (PDB 6CMI). The S239 residue is coloured red and ephrinB2 binding site is shown in purple. Left inset depict native S239 residue and its interaction with m102.3 HC (green) paratope. Right inset shows modelled L239 mutation. L239 was modelled using ChimeraX based on highest rotamer prevalence according to Dunbrack rotamer library. Below, the DS90-bound NiV F structure is shown, with S50 coloured in red. The left inset shows S50 interaction with DS90 CDR3. The right inset shows the I50 mutation, modelled using ChimeraX with the rotamer selected based on best agreement with the density map. (D) Neutralization (IC_50_) comparison of antibodies against NiV pseudoviruses with RBP mutations. Bar graphs show mean of duplicate IC_50_ values with standard deviation obtained from two independent assays run in triplicate. (E) ELISA K_D_ values (nM) of antibodies binding to prefusion NiV F or NiV F-S50I. Data expressed as mean of duplicate values with standard error obtained from ELISA curves in Figure S11C.

NiV passaged in sub-neutralizing concentrations of DS90 Fc (1 μg/mL for P1 & 2 and 2 μg/mL for P3) acquired a S50I escape mutation in F, which was almost completely fixed at 93% of reads (Figure 5B). The S50 residue resides within the DS90 epitope and sits adjacent to V101 of DS90 CDR3, which makes two backbone mediated hydrogen bonds with the side chain of N51 (Figure 5C). The introduction of the bulkier and hydrophobic isoleucine side chain in place of the native serine likely perturbs this bond network and allows escape from binding. To validate this, we recombinantly expressed NiV F with the S50I mutation and assessed binding to a panel of mAbs by ELISA. We found that DS90 Fc binding to NiV F-S50I was reduced ∼50-fold compared to wild-type prefusion F (Figure 5E, Figure S11E). In contrast, mAbs such as 5B3 and 12B2, which do not engage with the S50 residue, were unaffected and retained binding to the NiV F-S50I mutant. Interestingly, we were unable to produce NiV pseudoviruses with the F-S50I mutation to validate escape from DS90 Fc dimer. We hypothesise that this may be due to a low fusogenic ability of the F-S50I mutant, based on previous research which demonstrated that mutation of a nearby residue (L53D) caused reduced cell-to-cell fusion ^30^. To test this, we conduced cell-cell fusion assays comparing the F-S50I mutant to wild-type F (Figure S11F). Indeed, we observed significant reduction in fusogenicity of the S50I relative to wild-type, despite confirming similar levels of expression in 293T cells (Figure S11G).

This escape mutant experimental process was repeated for DS90-m102.4. Given lack of CPE in cell cultures at matched concentrations (Figure S11A), reduced concentrations were used for DS90-m102.4 starting at 0.2 μg/mL for P1 & 2 and increasing to 0.5 μg/mL for P3. Nanopore sequencing of P3 revealed no high prevalence of non-synonymous mutations and a genetic profile similar to that of NiV passaged in the presence of an isotype control mAb, indicating accumulation of only stochastic mutations within the genome (Figure 5B, Figure S11D). This was in agreement with neutralization assessment, where NiV passaged in DS90-m102.4 was still permissible to m102.4, DS90 Fc and DS90-m102.4 neutralization (Figure S11C).

### DS90-m102.4 offers protection against lethal NiV challenge

To assess the prophylactic efficacy of DS90 and DS90-m102.4 against NiV disease, the hamster model of infection was used. This model was previously established and has been used to assess the protective efficacy of RBP-specific mAbs^25,39,40^. Here, groups of hamsters (n = 6) were treated with antibody constructs (10 mg/kg) via intraperitoneal injection 1 day prior to infection with NiV-M (1000 LD_50_) (Figure 6A). Infected hamsters were monitored for 28 days post-infection. In the study design, 5B3 and m102.4 mAbs were included as benchmarks, as they were previously shown to be protective in ferrets against NiV disease^9,19^. As prophylaxes, DS90 Fc and DS90-m102.4 offered 100% protection from NiV disease and significantly improved survival over the non-specific control group, where all hamsters succumbed to infection by day 6 (Figure 6A). DS90 FcM did not improve the survival of hamsters compared to the control mAb, with all animals succumbing to infection by day 7. Interestingly, the 5B3 and m102.4 benchmarks offered 83% protection as prophylaxes, significantly improving survival over the non-specific control group. All surviving animals displayed weight gain throughout the experiment.

**Figure 6.**
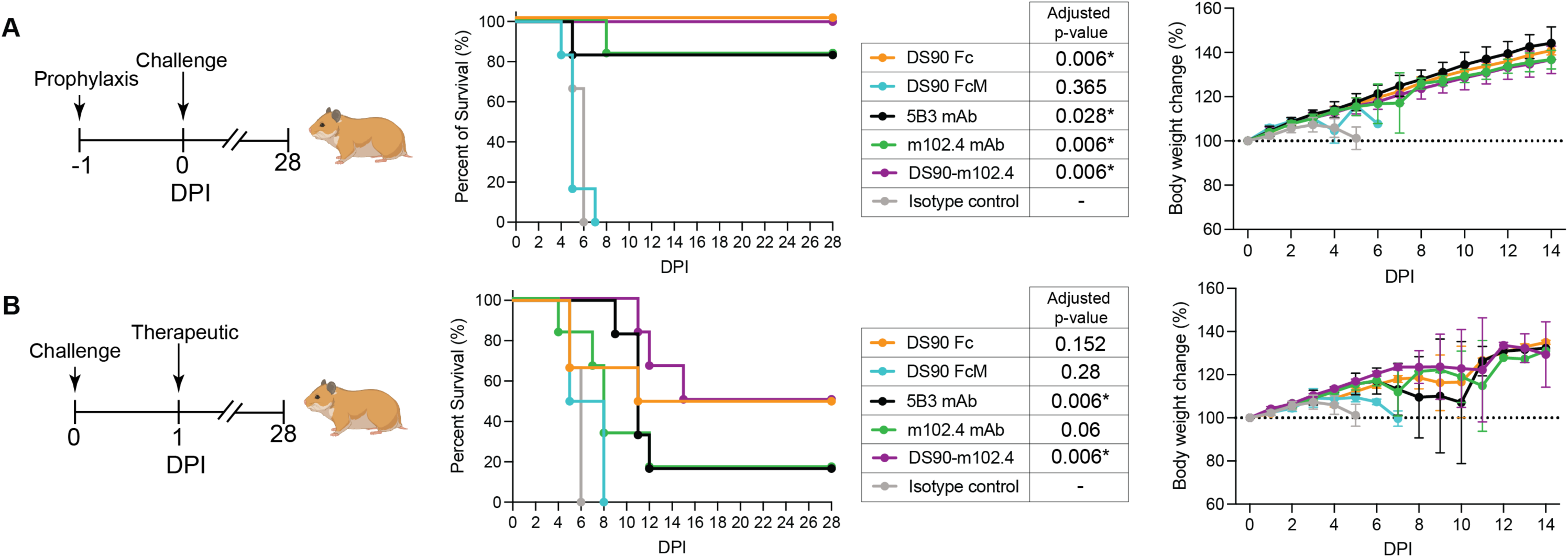
DS90-m102.4 and DS90 dimer offer protection against NiV-M in hamster model of infection. A hamster prophylaxis (A) or therapeutic (B) NiV-M challenge model. Six hamsters were treated with 10 mg/kg of antibody 1 day prior to (A) or after (B) infection via intraperitoneal injection with 1000 LD_50_ NiV-M. Survival curves are shown with percent body weight changes to the right. Statistics were conducted as multiple comparisons of treatment vs isotype control (C05 mAb) using log-rank (Mantel-Cox) test. P-values were adjusted for multiple comparisons using Holm-Šídák correction where α = 0.05. * indicates significant difference to non-specific isotype control.

Next, we assessed the post-exposure efficacy of DS90 and DS90-m102.4 using a similar study design. Here, groups of hamsters (n = 6) were infected with NiV-M (1000 LD_50_) and then treated with antibody constructs (10 mg/kg) 1 day after infection via intraperitoneal injection (Figure 6B). Interestingly, both DS90 Fc dimer and DS90-m102.4 therapeutics offered 50% protection against NiV-M disease, with DS90-m102.4 delaying mortality by an additional 4 days and providing significant improvement in survivability over the control group. DS90 FcM did not offer protection against infection, with all animals succumbing to infection by day 8. This is congruent with previous data showing markedly reduced neutralization by the DS90 monomer compared to the dimeric form (Figure 1C). Previously, m102.4 and 5B3 were shown to offer 100% protection from NiV disease as post-exposure therapy in ferrets. Interestingly, we found that as a therapeutic in hamsters, 5B3 and m102.4 only offered ∼17% survivability (1/6). This is in comparison to previous studies in ferrets, where therapeutic treatment of 5B3 offered a higher level of protection from NiV challenge, however multiple doses were given at significantly higher concentrations (∼25 mg/kg)^9,41^. Moreover, m102.4 was recently shown to only offer 17% survivability as a post-exposure therapy in non-human primates when administered 5 days post-infection at a higher dose of 25 mg/kg^26^.

### Humanization of DS90

Alpaca-derived nanobodies possess high sequence homology to human IgG heavy chain genes (75-90%)^42,43^. However, the efficacy of biologics of animal origin may be compromised in humans due to a reduced half-life *in vivo*. As such, we sought to humanize DS90 to enable future therapeutic use in humans. Sequence analysis revealed IGHV3-23*04 to be the closest human allele to DS90. To limit the effects of humanization on antibody efficacy, only residues outside of the CDRs were altered (Figure 7A). These include the following substitutions within DS90: A13P, R26F, G34S, Y36V, A48S, A74S, K86R, P87A, D88E and I92V. Humanized DS90 (hDS90) was expressed as a monomer (hDS90 FcM) in order to differentiate subtle changes in both ELISA affinity and NiV pseudovirus neutralization efficacy. Initially, we found that hDS90 FcM had a significant reduction in ELISA affinity to prefusion NiV F, also resulting in reduced neutralization in NiV pseudovirus assays (Figure 7B). To address this, we made use of the previously determined cryoEM structure to independently revert three FR residues back to the wild-type DS90 sequence that contribute to binding (V36Y, L78V, V92I) in separate constructs. Interestingly, hDS90 V36Y FcM ELISA binding to prefusion NiV F was improved 3.8-fold over that of wild-type DS90 FcM (hDS90 V36Y ELISA K_D_ 1.28 nM compared with wild-type DS90 ELISA K_D_ 4.92 nM). Subsequent pseudovirus neutralization tests revealed that hDS90 V36Y FcM offered similar neutralization levels to that of wild-type DS90 FcM (hDS90 V36Y IC_50_ 6.85 nM compared with wild-type DS90 IC_50_ 8.76 nM) (Figure 7B). Together, these results highlight the important contribution of FR residues to the nanobody paratope and therefore require consideration during the humanization process.

**Figure 7.**
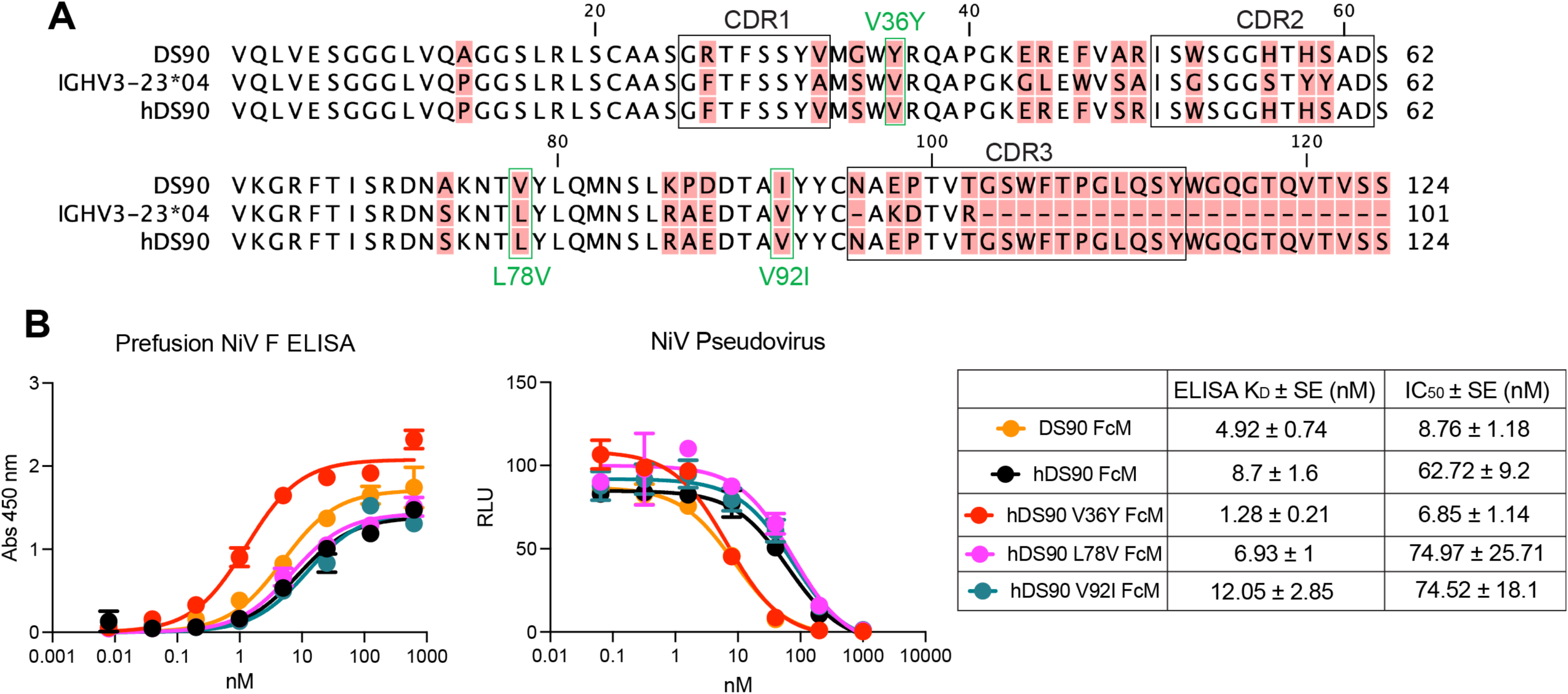
Design and testing of a humanised DS90 construct. (A) Sequence alignment of DS90 with IGHV-23*04 allele and hDS90. Black boxes represent CDR1-3 sequences. Green boxes represent selected reversions of hDS90 to wild-type sequence. Testing of hDS90 monomers for binding by ELISA (B) and neutralization by NiV pseudovirus neutralization tests (C). Data represents mean of triplicate values with standard deviation. NiV pseudovirus RLU is normalised to virus only wells. A summary of ELISA affinities and IC_50_ is shown to the right with standard error.

## Discussion

HNV continue to be a threat to global human health, with annual outbreaks of NiV in south-east Asia and emergence of novel HNVs globally^3,44–48^. Conventional antibodies present as promising therapies against HNV infection and disease, yet their efficacy may be compromised by viral escape^16,17,36^. Furthermore, the approved HeV equine vaccine and leading human vaccine candidates are based on RBP, which could potentiate the emergence of viral escape mutants within RBP, similar to what has been observed for SARS-CoV-2 RBD. Therefore, investigation of the F protein as a target for vaccines and therapeutics is warranted.

In this work, we described a first-in-class nanobody, DS90, that targets the prefusion form of both NiV and HeV F glycoproteins. We demonstrated its ability to cross-neutralize both authentic NiV and HeV with high potency and offer protection against NiV infection in a hamster challenge model. Through cryoEM, we show that DS90 binds a novel quaternary pocket, engages multiple domains and spans two F protomers. Together with functional assays, our data suggests that DS90 neutralizes NiV by stabilising the prefusion conformation of F, a mechanism utilised by other previously described mAbs such as 5B3^16^. In addition to this, we found that DS90 causes a local shift in the fusion peptide region of F and contacts residues previously shown to be important for F triggering by the RBP^30^. As such, DS90 may not only stabilise the prefusion form but may also interfere with triggering by RBP and prevent viral fusion and entry. Interestingly, we found that DS90 dimers exhibit much higher neutralization and protective effects in comparison to the monomeric form. Together, this data speaks toward the mechanisms underlying DS90 neutralization and protection. While we have not empirically determined whether this multivalent interaction occurs within one F trimer or between two adjacent prefusion F trimers, the distance between the C-terminus of the DS90 VHH domains is on the order of 8 nm within one trimer, which is incompatible with the flexible linker region between the VHH and Fc domains. Thus, the potency of DS90 may be governed by the ability to crosslink multiple prefusion F trimers on the surface of the virion, analogous to RBP mediated stabilization of F, and in turn may disrupt the higher order oligomers of F trimers that are required to coordinate viral and host cell membrane fusion^30^. This mode of action is also consistent with the finding that the CDR3 of DS90 assumes a similar binding mode to the adjacent F fusion loop within the dimer of trimer assembly, previously identified in both crystallographic and cryoEM studies (Fig S6D) ^30,31^. In this context DS90 dimers may mimic the native inter-prefusion F trimer interactions through bivalent engagement of adjacent F trimers and therefore stabilize and lock the F protein assemblies in the prefusion F conformation across the virion surface. As such, the F epitope occupied by DS90 within prefusion F may be targeted in HNV vaccines, small-molecule drugs designs and human mAbs, to yield potent neutralisation of HNVs.

NiV and HeV are RNA viruses that may readily mutate in the face of vaccine- or antibody-induced selection pressures. The ability of RNA viruses to mutate is best shown for SARS-CoV-2, where the omicron variant was able to escape both ancestral vaccine antibody responses as well as the majority of monoclonal antibodies in clinical development^20,49^. In the context of HNVs, three independent studies have trialled either bispecific strategies or antibody cocktails to limit NiV escape mutants^17,25,50^. However, these studies only investigated antibodies targeting epitopes within a single target (either F or RBP). In this work, we combined DS90 with the leading immunotherapy for NiV and HeV, m102.4, to create a bispecific antibody that is able to simultaneously target both F and RBP. We demonstrated that DS90-m102.4 is able to potently neutralize authentic NiV and HeV and improves upon both DS90 Fc dimer neutralisation of NiV-B04 and m102.4 neutralization of HeV.

In respect to viral escape, we showed that DS90-m102.4 was able to limit the emergence of escape mutations *in vitro*, demonstrating an advantage over the parental antibodies. This carries implications for therapeutic designs and administration in an outbreak setting, where repeated use of a single therapy may result in variants that are resistant to neutralization. We engineered the bispecific antibody with a short linker (G_2_SG_2_) between the DS90 VHH and m102.4. Given recent research describing molecular reach^51^; that is, the maximum distance between antigens enabling bivalent binding of an antibody, and its correlation with neutralisation potency, extending the linker region of DS90-m102.4 may allow for better neutralisation of NiV and HeV.

Results from this work also provide proof-of-concept for use of nanobody therapeutics, such as DS90, against NiV disease. Due to their simple domain architecture and lack of glycosylation sites, nanobodies can be efficiently expressed in prokaryotes, allowing for economical, expeditated production, and their high solubility and stability facilitates drug scaling, transportation and distribution - all of which are critical in response to viral outbreaks^52,53^. These excellent physiochemical properties also allow the option of nanobody pulmonary delivery through aerosolization, allowing for a targeted approach to combat respiratory infections^54,55^. Furthermore, due to its thermal stability, DS90 could additionally be used in rapid antigen tests in remote areas where access to refrigeration may be limited.

Despite their advantages, nanobodies are still vulnerable to escape mutations. Indeed, three passages of NiV in DS90 resulted in escape from neutralization, offered by a single amino acid substitution event (F-S50I) within the DS90 epitope. While this mutation ablates DS90 binding, our work corroborates previous experiments demonstrating that substitutions at this site result in reduce fusogenicity of F^30^. As such, it is possible that while NiV F-S50I may escape DS90 neutralising, this may come at a significant cost to viral fitness. Nevertheless, such escape mutations may be overcome by genetically linking DS90 to a RBP-specific nanobody, which may mirror the potency and resistance to viral escape as demonstrated by the DS90-m102.4 bispecific antibody. Alternatively, multiple nanobody domains that target divergent HNVs may be linked to produce a broadly neutralizing therapeutic against NiV, HeV and recently emergent LayV, to which current antibody therapies have no binding capacity^56^.

In summary, in this work we described a first-in-class nanobody therapy against both NiV and HeV, and humanised the sequence to allow for clinical translation. We also provided proof-of-concept demonstrating that bispecific immunotherapies that target both F and RBP limit the emergence of NiV escape mutations. This may carry implications into the NiV therapeutic field, where bispecific strategies or antibody cocktails may serve as improved therapies against disease and limit the emergence of variants of concern.

## Acknowledgements

The authors would like to acknowledge Dr Bill Close, Dr Matthias Flotenmeyer and Dr Lou Brillault and the facilities, and the scientific and technical assistance of the Australian Microscopy & Microanalysis Research Facility at the Centre for Microscopy and Microanalysis, The University of Queensland. We would like to thank Ian Mortimer (ITS Infrastructure Operations, UQ) for IT support. We would also like to acknowledge the Electron Microscopy Unit (UME) and technical assistant of Ximena Verges S, the Agricultural experiment station and the assistance of the alpacas herder Omer navarrete bahamonde. We thank the Centre for Biosafety Mega-Science, Chinese Academy of Sciences and the National Virus Resource Centre for resource support. A.I. would also like to thank Pedro the alpaca for the highly potent nanobodies that made this work possible.

## Author Contributions

Conceptualization: A.I., G.V.N., A.R.F., E.P., D.W. Methodology: A.I., G.V.N., X.Z., N.M., J.B., N.T., J.B.B., G.M., R.H.P., D.B., S.C., Y.S., D.W., A.R.F. Formal Analysis: A.I., G.V.N., X.Z., N.M., J.B., N.T., Y.S.L., Y.Y., D.P., J.B.B., R.H.P., S.C., A.R.F., D.W. Investigation & Validation: A.I., G.V.N., X.Z., N.M., J.B., G.M., N.T., Y.S.L., R.H.P., J.B.B., R.J., J.H., C.D., B.B.G., C.S.R., E.P., A.R.F. Resources: all authors contributed resources to this project.Writing – Original Draft: A.I., A.R.F., D.W. Writing – Review & Editing: all authors. Supervision: D.W., A.R.F., K.C. Y.S., S.C., G.M., D.B. Funding Acquisition: A.I., S.C., D.W., A.R.F.

## Figure Legends

**Figure S1 Biochemical characterisation of NiV F specific nanobodies.** (A) A representative image from Ficoll gradient separation of bacteria displaying nanobodies. Inset show scanning electron microscopy images of nanobody-expressing bacteria bound to beads coated with NiV F. (B) High resolution confocal microscopy of screened *E. coli* nanobody crude extracts against HeLa cells expressing NiV F under IRES-GFP. Scale bar shown is 20 μm (C) Sequence alignment of NiV F-specific nanobodies with red depicting conserved residues and blue depicting divergent residues SDS-PAGE (reducing and non-reducing) and ELISA of dimeric Fc (D) and monomeric FcM (E) nanobodies. (F) SDS-PAGE of bacterially expressed DS90-cMyc elution fractions under reducing conditions. (G) Thermal stability ELISA of bacterially-expressed and purified DS90 reactivity to NiV F. DS90-Myc (at a concentration of 10 nM) was incubated at a range of temperatures for 30 mins before adding as a primary antibody in an ELISA against NiV F. Data shown is an average of quadruplicate values with standard deviation and is normalised using mean RT value as 100%. (H) High resolution microscopy of bacterially-expressed and purified DS90-cMyc binding to HeLa cells expressing NiV F under IRES-GFP. Scale bar shown is 20 μm or 50 μm (zoom).

**Figure S2 Pseudovirus neutralisation assays of F-specific mAbs**. F-specific mAb panel was tested against NiV-M (A) and NiV-B08 (B) pseudoviruses to benchmark DS90 Fc dimer neutralisation. Data shown is an average of triplicate values with standard deviation. Calculated IC_50_ values in nM against both NiV-M and NiV-B08 are shown in (C). (D) Live virus neutralisation assay against NiV-B04 strain of DS90 Fc dimer compared with 5B3 mAb. Data shown is average of triplicate with standard deviation.

**Figure S3 Characterisation of the DS90 epitope.** (A) Prefusion NiV F was complexed with DS90 and ran through size-exclusion chromatography on a Superdex S200 Increase 10/300 GL column. Data is normalised to maximal values within each dataset. (B) Size-excluded fractions corresponding to major SEC peaks were concentrated and ran on an SDS-PAGE under reducing conditions. (C) Sequence alignment of HNV F protein ectodomains annotated with F domains. Blue circles depict hydrogen bond contacts and red circles depict salt bridges made by DS90 across all domains except HRB. NiV-M/98 Genbank: AAM13405; NiV-B/04 Genbank: AAY43915-1; NiV-B/12 Genbank: QHR79135; NiV-I/19 QKV44013; NiV-C/03 Genbank: QDJ04462; HeV-g1 Genbank: AEB21197.

**Figure S4 CryoEM processing workflow for NiV F-DS90.** (A) Acquisition and data processing workflow on cryoSPARC including representative 2D class averages and a representative micrograph. Patch Motion correction and Patch CTF estimation were performed. The aligned/motion-corrected micrographs were curated based on resolution estimation. and subset of selected particles were rebalanced and used for ab initio for initial models imposed C3 symmetry. The models were then used as references to classify ‘good’ selected 2D particles from all micrographs using heterogenous refinements. The final 200,640 particles were then further refined using non-uniform refinement, global CTF and local CTF refinement. (B) Final cryoEM map of DS90-bound NiV F trimer. (C) Fourier Shell Correlation (FSC) of final map. Dotted lines show cut-offs at 0.5 and 0.143. (D) Local resolution map of DS90-bound NiV F, generated in cryoSPARC. (E) 3D angular distribution plot of final DS90-F structure.

**Table S1** CryoEM data collection, processing and model building details.

**Table S2** Intermolecular forces present between DS90 and NiV F including hydrogen bonds and salt bridge. Table shows both donor (D) and acceptor atoms and the distance between them, calculated using UCSF ChimeraX.

**Table S3** Van der Waals interaction between NiV F and DS90. Table shows atoms involved in VDW overlap and the distance between the atoms, calculated using UCSF ChimeraX.

**Figure S6 DS90 causes a shift in N424 relative to other known NiV F structures.** (A) Ribbon diagrams of known NiV F structures (PDB 5EVM, 6TYS, 7UPD) superimposed on DS90-bound NiV F, with DS90 coloured gold and NiV F trimer coloured teal. Inset shows DS90 CDR3 in stick form, with oxygen coloured red and nitrogen blue, inserting into the NiV F pocket. Arrow highlights shift in orientation of N424 side chain relative to other known structures. The oxygen of main carbonyl group in P108 forms a hydrogen bond (red dotted line) with N424. (B) CryoEM workflow of symmetry expansion and local refinement of DS90 and Nipah F interface. Final map resolution of 3.59 Å was achieved. (C) Zoom of DS90-F local refinement (left) and apo-F (right). Apo-F structure was previously determined by others (PDB 8DNG/EMD-27566). Models are shows as cartoon depiction with N424 side chain shown as stick form. Associated sharpened map densities are shown in transparent grey. (D) Dimer-of-trimer assembly of NiV F (PDB 8DO4) with a single monomer of each F trimer coloured in purple or green. Inset shows the fusion loop region that is sequestered by the dimer-of-trimer interface. DS90 CDR3 is shown in orange and mimics the fusion peptide loop shown in purple.

**Figure S7 Structural comparison of DS90 epitope with other known mAbs**. The epitopes of apical, lateral or basal Nipah F mAbs relative to the DS90 footprint coloured in orange on Nipah F surface.

**Figure S8 Sequence conservation between HNV F proteins.** (A) Surface representation of the DS90 footprint highlighted in blue on NiV F. NiV F is coloured by conservation according to sequence alignment in Figure S3. Residues that differ between NiV-M and NiV-B04 F proteins (A249 and D252) and NiV-M and HeV (T263) are annotated. (B) NiV pseudovirus mutant neutralisation by DS90 Fc dimers. Data shown is an average and standard deviation of 6 replicates, and IC_50_ and standard error calculated from two experiments conducted in triplicate.

**Figure S9 Biochemical characterisation of DS90-m102.4 and supporting data for ADCC assay.** (A) Coomassie stained, non-reducing SDS-PAGE of purified m102.4 mAb, DS90-m102.4 and DS90 nanobody. (B) Size-exclusion chromatography of m102.4 IgG and DS90-m102.4 ran on Superdex S200 Increase 10/300 GL column. Data is normalised to maximal value within each dataset. (C) Indirect ELISA of mAbs against NiV RBP or prefusion NiV F. Data shown is average of triplicate data with standard deviation. (D) Authentic NiV neutralization by antibodies and nanobody Fc dimers conducted in BSL4 shown as cytopathic effect endpoint titre (CPE EPT). Data shown is an average of duplicate values.

**Figure S10 ADCC of NiV F and NiV RBP expressing cells.** (A) ADCC activation curves of antibody constructs targeting NiV F or NiV RBP. Data shown is average of triplicate values with standard deviation. (B) Validation of RBP and F expression on 293T cells by flow cytometry with 5B3 (F-specific) and m102.4 (RBP-specific) mAbs. Non-specific antibody (C05) used as a negative isotype control.

**Figure S11 Escape mutation analysis of DS90-m102.4.** (A) Cytopathic effect (CPE) monitoring of Vero cells inoculated with authentic NiV-M in the presence of antibody at 72 hrs post infection. Mock and virus only controls included at 24 hrs post infection (B) CPE monitoring of Vero cells infected with authentic NiV-M in the presence of DS90-m102.4 at 0.1 μg/mL and 0.2 μg/mL. The cell monolayer for DS90-m102.4 at 0.1 μg/mL at day 5 post infection was completely ablated and no image was acquired. (C) Neutralization analysis of authentic NiV viral stocks passaged in antibody constructs from P3. Virus escape is seen for m102.4 and DS90 but not DS90-m102.4. (D) Incidence of escape mutations generated by antibody selection pressures within the F-RBP cDNA amplicon. Black dots represent synonymous mutations, grey circles represent stochastic mutations generated by a non-specific antibody (C05), red circles represent non-synonymous mutations. Data shown are additional replicates conducted for m102.4 and DS90-m102.4. (E) ELISA curves of antibodies binding to recombinantly expressed prefusion NiV F or NiV F-S50I. Data shown is mean of duplicate values with standard deviation. (F) Cell-to-cell split reporter luciferase fusogenicity assay of wild-type NiV F and NiV F-S50I mutant in 293T cells. Statistical analysis conducted was Tukey’s multiple comparison one-way ANOVA. **** indicates p < 0.0001. (G) Expression levels of NiV F and NiV F-S50I in 293T cells, quantified by flow cytometry and 12B2 staining. C05 used as a non-specific isotype control.

## Materials & Methods

### Cell lines

HEK293T cells (ATCC) were maintained in DMEM supplemented with 1 mM sodium pyruvate, penicillin (100 IU/mL) streptomycin (100 μg/mL), and 10% heat-inactivated (HI) fetal calf serum (FCS, Bovogen). BHK-21 cells (ATCC) were maintained in DMEM supplemented with penicillin (100 IU/mL) streptomycin (100 μg/mL) and 5% HI-FCS. Vero cells (ATCC) were grown in MEM (Cat# 11095080, Gibco, Thermo Fisher Scientific) supplemented with 10 mM HEPES (Gibco, Thermo Fisher Scientific), 1% (v/v) antibiotic– antimycotic (Cat#15240112, Gibco, Thermo Fisher) and 10% (v/v) FBS (CellSera Australia, NSW). Cells were cultured at 37 °C and 5% CO_2_. ExpiCHO cells (ThermoFisher) were maintained in ExpiCHO Expression media as per manufacturer’s instructions. All cell lines were tested for mycoplasma and found to be mycoplasma-free (MycoAlert Mycoplasma Detection Kit, Lonza).

### Viruses

Nipah Malaysia/Human/1999 isolated from the cerebrospinal fluid of an infected human was passaged and grown in Vero cells to create a working stock with a titre of 1.89 x 10^7^ TCID_50_/ml and used for escape mutant and neutralization experiments at CSIRO ACDP. HeV/Australia/Horse/1994/Hendra was originally isolated from the lung of a horse and propagated in Vero cells for use in *in vitro* neutralization assays. The Nipah virus Bangladesh strain (CSTR: 16533.06.IVCAS 6.7488) used for neutralisation assays and Malaysia strain (CSTR: 16533.06.IVCAS 6.7489) for hamster challenge were acquired from the National Virus Resource Centre at the Wuhan Institute of Virology, Chinese Academy of Science.

### Animal models

For immunization a seven years old, male alpaca was used, kept at the experimental station of the Santa Rosa Ranch, University Austral of Chile. The whole process followed the guidelines established by the Bioethics Committee of the Austral University of Chile (certifications 338/2019 and 388/2020).

Female Syrian golden hamsters, aged five to six weeks, were obtained from Wuhan Biological Products Institute and housed in SPF animal facilities at the Wuhan Institute of Virology, Chinese Academy of Science. Viral infection was conducted in a ABSL-4 facility in accordance with the guidelines for the care and use of laboratory animals and the Institutional Review Board of the Wuhan Institute of Virology, Chinese Academy of Science.

### Antigen production

The NiV Malaysian strain F ectodomain (1-483, NCBI: NP_112026.1) stabilized in the prefusion conformation by the T4 fibritin foldon domain was produced in the ExpiCHO system. ExpiCHO cells (Thermo Fisher) were transfected as per manufacturer’s instructions, and supernatant containing secreted proteins was harvested 7 days post transfection. NiV F was purified using an in-house made HiTrap NHS column (Cytiva) conjugated with ectodomain-specific mAb66^15^, followed by extensive washing with PBS and elution with high pH buffer (100 mM Tris-HCl, 400 mM NaCl, 5 mM EDTA, 20 mM DEA, pH 11.5). Elution fractions were neutralized with 1 M Tris-HCl pH 6.8 before buffer exchange into PBS using a 30 kDa MWCO spin concentrator (Merck Amicon).

The NiV Malaysian strain RBP (71-602, GenBank: NP_112027.1) was expressed with a N-terminal mouse Ig heavy signal sequence (MGWSCIILFLVATATGVHS) and a C-terminal c-Tag (EPEA) linked by a GSG linker in the ExpiCHO system (ThermoFisher). Cell transfection and supernatant harvest was conducted as described for NiV F. NiV RBP was purified from cell supernatant using CaptureSelect^TM^ C-tag Affinity column (ThermoFisher) with a Tris-buffered saline (TBS, 100 mM Trizma Base, 150 mM NaCl, pH 7.4) wash buffer and a high salt elution buffer (20 mM Trizma Base, 2 M MgCl_2_, pH 7.4). Elution fractions were collected and buffer exchanged to TBS using a 30 kDa MWCO spin concentrator (Merck Amicon).

### Antibody production

Antibody sequences were acquired from published sources^7,15–17,24,57^ and cloned into expression vectors as previously described^32,58^. The bispecific antibody HC was generated by overlap PCR of the DS90 VHH domain to the N-terminus of the m102.4 HC, joined by a G2SG2 linker. The PCR product was subcloned into an antibody expression vector as previously described ^58^. All antibodies were expressed in the ExpiCHO system. Briefly, a ratio of 2:1 LC to HC plasmid DNA was used to transiently transfect ExpiCHO cells as per manufacturer’s guidelines (Thermo Fisher). Supernatant containing secreted antibodies was harvested 7 days post transfection. Antibodies were purified from supernatants using a protein A column (Cytiva), washing with a wash buffer (25 mM Tris, 25 mM NaCl, pH 7.4) and eluting antibodies with a low pH buffer (100 mM sodium citrate, 150 mM NaCl, pH 3). Eluted fractions were neutralized with 1.5 M Tris-HCl pH 8.8 and buffer exchanged to PBS using a 30 kDa MWCO spin concentrator (Merck Amicon).

### Alpaca immunisations

Alpaca immunizations were conducted as previously described^29^, and were adhered to the guidelines established by the Bioethics Committee of the Austral University of Chile (certifications 338/2019 and 388/2020). A pre-immune blood sample of 5 mL was acquired one day before immunization for a pre-immune serum control. On the immunization day (day 1), 200 μg of the NiV F protein was resuspended in 2 mL of sterile water and further dissolved in 2 mL of adjuvant (Veterinary Vaccine Adjuvant, GERBU FAMA), then subcutaneously immunized into a male alpaca (*Vicugna pacos*). A total volume of 4 mL was administered at four different locations on the alpaca. Immunization was repeated at day 14, 28 and 42. On day 46 after the first immunisation, 120 mL blood sample was collected from the jugular vein in tubes containing 3.8% sodium citrate as an anti-coagulant. The uncoagulated blood sample was mixed with an equal volume of HBSS medium without calcium (Gibco), divided into 10 mL aliquots, and carefully layered on top of 5 mL of Ficoll-Paque Premium (GE Healthcare) in 15 mL sterile Falcon tubes. Following centrifugation (1200 rpm, 80 min at room temperature), the PBMC fraction was recovered from the interphase, washed twice in PBS 1X by centrifugation at 3500 rpm for 10 min, and then resuspended in 4 mL of sterile PBS 1X (Gibco). RNA extraction and cDNA production were carried out using the commercial RNeasy Mini Kit (Qiagen) and QuantiTect Reverse Transcription Kit (Qiagen), respectively, following the manufacturer’s instructions.

### Bacterial nanobody display library construction

2 μL of cDNA were used as templates in 50 μL PCR reactions with oligonucleotides CALL001 (5’-GTC CTG GCT CTC TTC TAC AAG G-3’) and CALL002 (5’-GGT ACG TGC TGT TGA ACT GTTCC-3’)^59^. The resulting amplified fragments, ∼700bp for VHH-CH2 domains and ∼1 kb for conventional VH-CH1-CH2 domains, were separated on 1.2% (w/v) low melting agarose gels. The ∼700bp band was subsequently extracted and purified using the QIAEX II Gel Extraction kit (Qiagen). The purified fragment served as a template for a second PCR reaction with oligonucleotides VHH-Sfi2 (5’-GTC CTC GCA ACT GCG GCC CAG CCG GCC ATG GCT CAG GTG CAG CTG GTG GA-3′) and VHH-Not2 (5’-GGA CTA GTG CGG CCG CTG AGG AGA CGG TGA CCT GGG T-3’) to ultimately obtain amplified fragments of ∼400 bp, corresponding to VHH domains. The amplified VHH fragments were digested with *Sfi*I and *Not*I restriction enzymes and then ligated into the same restriction sites of the vector pNeae2^60^.

The ligations underwent electroporation in *E. coli* DH10B-T1 R cells. This size was determined by plating on LB-chloramphenicol agar plates with 2% w/v glucose, incubated at 37 °C. Re-ligated vectors were estimated from a control ligation performed concurrently without the DNA insert. The transformed bacteria were scraped from plates and stored at −80 °C in LB broth with 30% glycerol.

### Density gradient separation

200 μg of NiV F recombinant protein in PBS was coupled to NHS-activated Sepharose 4 Fast Flow beads. 300 μL of NiV F beads were incubated with 2 mL of the induced bacterial display library (chloramphenicol resistant) mixed with a negative control bacteria (ampicillin resistant) in a 1:1 ratio. Incubation was performed in 15 mL conical low binding tube on a rocking platform for 30 minutes at room temperature.

The mixture was gently added on top of 6 mL of Ficoll (Ficoll-Paque PLUS, GE Healthcare) in a 15 mL conical tube and centrifuged at 200×g for 1 minute. The unbound fraction (upper fractions) was carefully discarded, leaving a visible pellet of beads that was resuspended in 4 mL PBS and rotated for 5 minutes at room temperature. This was repeated six times. Finally, 1 mL of LB medium was added and incubated for 5 minutes at room temperature. Subsequently, 50 μL of the mixture was plated on LB agar plates with 50 μg/mL ampicillin and 2% glucose. An additional 50 μL was plated on LB agar plates with 25 μg/mL chloramphenicol and 2% glucose, and the remainder on at least two LB chloramphenicol/glucose agar plates. All plates were incubated at 37 °C overnight. The colony count from the first two plates was used as a metric for the specific enrichment of bacteria expressing nanobodies from the library^29^.

### Immunofluorescence assay

Individual colonies obtained from a density gradient separation protocol were inoculated into 2 mL of LB medium and incubated overnight at 37 °C with agitation at 200 rpm. 100 μL of pre-inoculum was added to 1.9 mL of fresh LB medium with 25 μg/mL chloramphenicol, incubated at 37 °C with 200 rpm agitation until it reached OD_600_ of 0.45–0.6. To induce protein expression, IPTG was added at a final concentration of 50 μM for 3 hr at 30 °C and 200 rpm. The culture was pelleted and resuspended in 1 mL PBS with 0.2% TritonX100, sonicated for 10 s at 40% three times on ice, then centrifuged at 14,000 ×g for 30 min at 4 °C and the supernatant was recovered to obtain a total protein extract from each clone.

In parallel, HeLa cells were maintained at 37 °C in DMEM supplemented with 10% FCS and 100 units/mL of penicillin and streptomycin. Plasmid transfection was performed in 10 cm plates using 25 μg of NiV F-GFP plasmid using Lipofectamine 2000 (Invitrogen). Culture media was supplemented with Normocin (Invivogen). 24 hr after transfection, cells were split into 96 well optical plates (Thermo Fisher) at a density of ∼10,000 cells per well.

Cells were washed three times with PBS and fixed with 4% paraformaldehyde at room temperature for 30 minutes. After fixation, PBS washing was performed three times, followed by permeabilization in PBS containing 0.2% Triton X-100. The cells were washed again three times with PBS before being incubated with an extract containing nanobodies overnight at 4°C.

The following day, the cells were washed 3 times with PBS. A mouse anti-myc antibody (Cell Signalling) was then applied at a dilution of 1:3000 and incubated for 45 minutes at 37 °C. Subsequently, the cells were washed three times with PBS and incubated with an anti-mouse Alexa 647 secondary antibody and 0.1 mg/mL DAPI for 30 minutes at 37 °C. After the final three washes with PBS with 0.5% Tween20, fixed cell images were acquired using a high-content automatic microscope Celldiscoverer 7 (Carl Zeiss GmbH, Jena, Germany).

### Nanobody production

Nanobody sequences were cloned into a mammalian expression plasmid with a C-terminal monomeric human Fc tag (FcM) or dimeric Fc tag via In-Fusion cloning (Takara Bio). Constructs were transfected into ExpiCHO cells using ExpiFectamine as per manufacturer’s instructions. Supernatants were harvested on day 7 by centrifugation at 4800 ×g for 20 mins before filter sterilization. Harvested supernatants were then passed through a protein A column (Cytiva) before extensive washing with wash buffer (25 mM Tris, 25 mM NaCl, pH 7.4) and elution of nanobodies with a low pH buffer (100 mM sodium citrate, 150 mM NaCl, pH 3). Eluted fractions were neutralized with 1.5 M Tris-HCl pH 8.8 and buffer exchanged to PBS using a 30 kDa MWCO spin concentrator (Merck Amicon).

Nanobodies were also subcloned into pHen6, an *E. coli* periplasmic expression vector, using SfiI and NotI restriction sites. For periplasmic expression, WK6 bacteria were transformed with the pHen6-DS90 vector and further scaled into 1 L Terrific Broth (TB) medium containing 100 μg/ml ampicillin, 2 mM MgCl_2_ and 0.1% glucose, and incubated at 37 °C to an OD_600_ of 0.6–0.9. Nanobody expression was induced by adding 1 mM of IPTG for 20 hr at 28°C. The collected bacteria were resuspended in 12 mL TES buffer (0.2 M Tris pH 8.0, 0.5 mM EDTA, 0.5 M sucrose) and incubated for 1 hr at RT, then incubated for another hour with extra 18 mL TES/4 buffer (TES buffer diluted 4 times) for osmotic disruption. The lysate was centrifuged at 3260 ×g at 4°C, and nanobodies were purified from the supernatant by a Ni-NTA agarose purification.

### Surface plasmon resonance

Prefusion NiV F and NiV RBP head domain proteins were produced with an AviTag™ and biotinylated using BirA (Avidity), as per manufacturer’s instructions. Briefly, proteins were buffer exchanged into low salt PBS (20 mM NaCl). BirA protein was added to AviTagged proteins at 2.5 μg BirA per 10 nmol of AviTag™ protein with 1X Supermix buffer. Reaction was allowed to occur for 1 hr at room temperature. AviTag™ proteins were then purified by size-exclusion chromatography on a Superose 6 Increase column (Cytiva), with fractions of interest concentrated using 30K MWCO Amicon tubes.

SPR was performed using a Biacore 8K+ instrument (Cytiva). Biotinylated NiV F and NiV RBP head were diluted to 2.5 μg/mL in running buffer (HBS-P+, pH 7.4, Cytiva) and used to coat streptavidin (SA) series S sensor chips (Cytiva), as per manufacturer’s guidelines. Three-fold serial dilutions were included for monomeric nanobody constructs starting at 50 nM, dimeric nanobody constructs starting at 12.5 nM and antibodies starting at 6.7 nM. Antibodies were injected over both Fc1 and Fc2 of channels 2 and 3 in series for 180 sec at 30 μL/min, followed by a dissociation period of 1500 sec. The SA chip was regenerated after each cycle using 10 mM glycine pH 1.5. The data was double reference subtracted; reference cell subtraction (Fc1) from the active cell (Fc2), and zero concentration subtraction for each analyte-antibody pair.

Tandem SPR was similarly conducted. Briefly, antibody constructs were diluted to 50 nM and injected over captured biotinylated NiV F or NiV RBP head domain with a contact time of 180 sec, flow rate of 30 μL/min and dissociation time of 60 sec. This was followed by the second analyte (unbiotinylated NiV F or NiV RBP head) at a concentration of 100 nM, with a contact time of 180 sec, flow rate of 30 μL/min and dissociation time of 1200 sec. The SA chip was regenerated using 10 mM glycine pH 1.5.

All sensorgrams for the association and dissociation phases of monomeric constructs were fitted to a 1:1 binding model and bivalent constructs were fitted to bivalent analyte general kinetics model using Biacore Insight Evaluation software (Cytiva) and graphed using GraphPad Prism 10 software.

### ELISA

Nanobody binding was evaluated by ELISA to determine antigen specificity. Briefly, 2 μg/mL of antigen was coated on a Nunc MaxiSorp 96 well plate overnight at 4 °C. All wells were blocked with 150 μL of blocking buffer (5% KPL milk diluent solution (SeraCare) in PBS with 0.1% Tween20 (PBST)) for 1 hr at room temperature. Plates were then probed with a serial dilution of mAb or Fc-tagged nanobody for 1 hr at 37 °C. Plates were then washed thrice in water before addition of HRP-linked goat anti-human secondary (Sigma-Aldrich) diluted 1:2500 in block for 1 hr at 37 °C. Plates were washed as before and assay was revealed with TMB chromogen solution (Life Technologies) for five minutes before halting reaction with 1 M H_2_SO_4_. Absorbance was read at 450 nm and data was plotted with background binding subtracted on GraphPad Prism 9 using a one-site specific model.

Thermal stability ELISAs of bacterially-expressed DS90-cMyc were conducted as above with the following amendments. At the primary step, DS90-cMyc was incubated at a concentration of 10 nM at different temperatures (25−90 °C) for 10 mins and then cooled on ice before addition to the antigen-coated plate. At the secondary step, a mouse anti-cMyc antibody (Cell Signalling) was used diluted 1:3000 for 1 hr at 37 °C. After washing, a tertiary HRP-linked goat anti-mouse antibody (Sigma Aldrich) was diluted to 1:5000 in block and added to wells for 1 hr at 37 °C.

### Size-exclusion chromatography

Size-exclusion chromatography was used for either purification of DS90-complexed NiV F or size and homogeneity analysis of bispecific antibodies. Here, 50-350 μg of protein was loaded onto a Superdex 200 Increase 10/300 GL (Cytiva) column in 500 μL buffer. Peak 1 mL fractions were collected and concentrated using a 10 or 30K MWCO (0.5 mL) centrifugal filter units (Amicon Merck) for downstream analyses.

### CryoEM data collection and processing

DS90-bound F glycoproteins were purified by size-exclusion chromatography and concentrated to 0.04 mg/mL, before applying 2 μL of sample to grow-discharged Quantifoil 200 mesh copper grids (Q2/2). Grids were plunged frozen using EMGP2 Leica system and imaged on CRYOARM300 (JEOL) equipped with an in-column Omega energy filter and K3 detector (Gatan) in counting mode at the University of Queensland Centre for Microscopy and Microanalysis. Two datasets of movies (5971 and 3116) were acquired with 87.5 (e^-^/Å^2^) using SerialEM 3.3^61^ at magnification of 50,000× with a defocus range of -0.5 to -3.0 μm. All movies were motion-corrected by patch motion correction and processed by patch CTF estimation using cryoSPARC 3.3^62^. Next, template-free particles were picked and 2D classification was performed for template picking. The particles from template-based picking were sorted by 2D classification, *ab initio* reconstruction (C3 symmetry) and heterogeneous refinement (C3 symmetry). Selected particles were refined with C3 symmetry by non-uniform refinement and subsequent global and local refinements were performed. Further non-uniform refinement was used to refine the final map. Data processing details are shown in Figure S3 and Figure S4.

### CryoEM model building

An existing NiV F structure (PDB 5EVM) was used as an initial model^30^. Additional protein and glycan residues were added using Coot v.0.9.8.161. Refinement of the models was performed using ISOLDE v.1.362 and the quality of the model and fit to density was determined using Phenix v.1.20.1–448763. All figures were made using UCSF Chimera v.1.17 and ChimeraX v.1.364. Model generation and acquisition details for are presented in Figure S4 and Table S1.

### Micro-fusion inhibition tests

Cell-cell fusion assay was conducted as previously described^63^. HEK293T Lenti rLuc-GFP 1– 7 (effector cells) and BHK-21 Lenti rLuc-GFP 8–11 (target cells) were seeded separately at 7.5×10^5^ cells per well of a 6-well plate. Effector cells were transfected the following day with 500 ng pcDNA3.1 NiV F and pcDNA3.1 NiV G (Malaysia strain) or 250 ng pcDNA3.1 HeV F and pcDNA3.1 HeV G glycoproteins using the TransIT-X2 Dynamic Delivery System as per the manufacturer’s recommendations (Mirus). Dimers and monomers of nanobodies were diluted 2-fold, from a starting concentration of 50 μg/mL in a white-bottomed, sterile 96-well plate (Corning) at a final volume of 25 μL/well. No-nanobody controls were also included to determine cut-off values of inhibition. The nanobodies were incubated with 2×10^4^ effector cells in 50 μL for 1 hr, after which the equivalent number of target cells were co-cultured to corresponding wells and incubated for 18 hrs. To quantify Renilla luciferase expression media were replaced with 50 μL of diluted substrate, Coelenterazine-H, 1 mM (Promega) 1: 400 with PBS. The plate was incubated in the dark for 2 min then read on the GloMax Multi^+^ Detection System (Promega). Data was fitted with a four-parameter inhibitor vs response model with a top constraint of 100 and a bottom constraint of 0 on GraphPad Prism 10 software.

### NiV pseudovirus assays

The neutralization capacity of nanobody constructs was initially assessed via a Nipah lentivirus-based luciferase reporter system. NiV pseudoviruses were produced as previously described^32,63^. A day prior to transduction, 2 x 10^4^ target BHK-21 cells were plated on a white Nunc MicroWell 96-well plate in DMEM 5% FCS (D5) media. mAbs or nanobodies were serially diluted in D5 media and incubated with NiV pseudovirus for 1 hr at 37 °C with 5% CO_2_. Next, virus/inhibitor mixtures were transferred onto target cells and incubated for a further 72 hrs. Pseudovirus reporter activity was measured by discarding supernatant and adding 50 μl/well of a 1:1 mix of Bio-Glo Luciferase Assay System (Promega) and serum-free Opti-MEM for 10 mins. Luminescence was quantified using a VarioSkan LUX system (ThermoFisher).

### Generation of 293T stable cells expressing NiV glycoproteins

To generate stable target cell lines expressing NiV glycoproteins, HEK293T cells were cultured at 37°C 5% CO_2_ in DMEM (Gibco) with 10% Foetal Bovine Serum (SAFC Biosciences Bovogen Biologicals) before being transfected using the Lipofectamine 2000 system (Invitrogen) with the stable expression vector pNBF (National Biologics Facility, Queensland, Australia) encoding either full length NiV-F or NiV-RBP under a CMV promoter and a puromycin resistance gene. At 48 hours post-transfection, Puromycin (Gibco) was added to the cells at a concentration of 0.4 µg/ml to select for positive transfectants. For cell sorting, cells transfected with pNBF NiV-F or pNBF NiV-RBP were stained using either mAbs 5B3 or m102.4 at a concentration of 2 µg/ml, respectively. Sorted cells were propagated in Puromycin as above before successive cell sorting was performed until positive cell populations reached >90% NiV glycoprotein expression. Stable cell lines expressing NiV glycoproteins were subsequently used in ADCC assays or frozen and stored in liquid nitrogen for future use.

### Antibody-dependent cellular cytotoxicity assay

ADCC assays were performed using previously generated stable HEK293T target cell lines expressing NiV glycoproteins in combination with the ADCC Bioassay Effector Cells G7102 system (Promega) in accordance with the manufacturer’s guidelines. Briefly, target cell lines were seeded in the inner 60 wells of white 96 well plates (Corning) at a concentration of 2.5×10^4^ cells/well. The following day, 3-fold serial dilutions of mAbs and effector cells at an effector:target ratio of 6:1 were added. After 18 hours, Bio-Glo luciferase substrate (Promega) was added before luminescence was read using the FLUOstar Omega microplate reader system (BMG LabTech). ADCC curves and associated values were generated using GraphPad Prism.

### Authentic NiV & HeV neutralization assays

Standard virus neutralization tests (VNT) were performed in the Biosafety Level 4 (BSL4) high containment laboratory at the CSIRO Australian Centre for Disease Preparedness. Nanobody or antibody constructs were double diluted in duplicate in MEM media (Gibco) starting at a concentration of 200 µg/ml in 96-well tissue culture plates. An equal volume of 200 TCID_50_ of either NiV or HeV was added to the nanobodies or antibody constructs and incubated for 30 min at 37 °C in a 5% CO_2_ humidified incubator. After incubation, Vero cell suspension containing 2 x 10^5^ cells/ml prepared in MEM-10 cell media was added to all wells in the tissue culture plates and then incubated for 4 days. After 4 days, the wells were observed for the presence of viral CPE and the neutralizing concentrations were determined as the concentration where 100% virus neutralization was seen in duplicate wells.

Plaque reduction neutralization assays were also conducted for NiV-M and HeV at the CAS Key Laboratory of Special Pathogens and Biosafety facility under BSL4. Briefly, antibodies were serially diluted four-fold and incubated with a target of ∼100 plaque-forming units (PFU) of NiV-M or HeV for 60 min at 37 °C. Virus and antibody mixtures were then added to individual wells of six-well plates of VeroE6 cells. After an additional 1 h incubation at 37 °C, the antibody-virus mixture was removed, and DMEM containing 2.5% FBS and 0.9% carboxymethylcellulose was added. Plates were fixed with 8% paraformaldehyde and stained with 0.5% crystal violet and rinsed thoroughly with water 5 days later. Neutralization potency was calculated based on PFU for each virus in the well without antibody. Plaques were then enumerated, and the neutralization percentage was calculated by the formula: Neutralization (%) = (1 – sample plaques/virus control plaques) (%). The IC_50_ was determined using GraphPad Prism v10 software. The experiments were performed in triplicate with independent virus preparations and duplicate readings for each replicate.

### NiV escape mutant generation

NiV escape mutants resistant to neutralization were generated by incubating nanobody or antibody constructs diluted in 100 µl MEM at two different sub-neutralizing concentrations for each (10 µg/ml for C05, 2 µg/ml for m102.4, 1 µg/ml for DS90 and 0.2 µg for DS90-m102.4) with 100 µl MEM containing 1 x 10^5^ TCID_50_ NiV for 30 min at 37 °C in a 5% CO_2_ humidified incubator in the BSL4 lab. After incubation, the 200 µl nanobody-virus mix was inoculated onto 1 x 10^6^ Vero cell monolayers prepared in 6-well plates for 30 min at 37 °C in a 5 % CO_2_ humidified incubator with gentle rocking every 10 min. The wells were then topped up with 800 µl MEM-10 cell media and placed back into the incubator. The plates were checked daily for the presence of viral CPE and wells were harvested when CPE was present (2 dpi to 5 dpi). The virus containing supernatants were then passaged two more times using the same protocol and nanobody concentrations, except for the third passage of the DS90 nanobody and the DS90-m102.4 fusion nanobody, where 2 µg/ml and 0.5 µg/ml were used, respectively, to increase selection pressure.

For sequencing analysis, 50 µl of supernatant from each passage was aliquoted into 260 µl MagMAX™ lysis buffer and removed from BSL4 for RNA extraction using a MagMAX™-96 Viral RNA Isolation Kit (Applied Biosystems) on a KingFisher Flex (ThermoScientific). Samples were then precipitated with isopropanol, washed with 70% ethanol and then used for NGS sequencing.

### Escape mutant deep sequencing and bioinformatic analysis

To identify escape mutations in the F and G genes of NiV, Nanopore sequencing of viral amplicons was conducted. Extracted RNA from cell culture supernatant was generated into cDNA and PCR amplified using SuperScript™ III RT/Platinum™ Taq Mix with 100 ng of RNA per sample with the following primer set: NipFG_F 5’-TTTCTGTTGGTGCTGATATTGCGGAGACCTTCTAACAGCCAGG-3’ and NipFG_R 5’-ACTTGCCTGTCGCTCTATCTTCCGGGCCGAAAATACTAAGTCTC-3’. The cDNA synthesis and pre-denaturation conditions were 50 °C for 30 minutes and 94 °C for 2 minutes, respectively. PCR amplification was performed under the following conditions: 94 °C for 15 seconds (denaturation), 55 °C for 30 seconds (annealing), and 68 °C for 4 minutes and 30 seconds (extension) for 30 cycles with a final extension at 68 °C for 5 minutes.

Following amplification, PCR fragments were gel-purified using the Monarch® DNA Gel Extraction Kit (T1020, New England Biolabs, USA) and barcoded using the PCR Barcode Expansion kit (PBC001, Oxford Nanopore, UK). The barcoded fragments were then gel-purified, pooled in equimolar ratios, and prepared for sequencing using the Ligation Sequencing Kit (SQK-LSK109, Oxford Nanopore, UK). Libraries were quantified with the Qubit™ dsDNA HS Assay Kit (Thermo Fisher Scientific, USA), and 50 fmol of the final library was loaded onto a Nanopore Flowcell (FLO-MIN106D, Oxford Nanopore, UK).

Base-calling and adapter trimming were executed using guppy_basecaller (Version: 3.3.0+ef22818). The base-called reads in fastq format were aligned to the NiV NV/MY/99/VRI-0626 genome (Genbank: AJ627196) using the Burrows–Wheeler Aligner with the flags mem -x ont2d^64^. The resulting BAM files were filtered for quality, retaining only reads with a minimum length of 500 bp and mapping quality >20. The depth of coverage was calculated using samtools (v1.3). Per base sequence variation and amino acid changes were identified using iVar (v1.4.2)^65^ with a minimum quality score threshold of 20 and a minimum frequency of 0.03. Variant frequencies and alignment depth were visualised using GraphPad Prism (v10.0.2), and individual variants were validated using Integrative Genomics Viewer (v 2.16.0). All fastq data generated in this study have been deposited in the NCBI Sequence Read Archive with accession number PRJNA1022870.

### NiV hamster challenge

Female Syrian hamsters (five-to-six weeks of age) were purchased from Wuhan Institute of Biological Products Co., Ltd. To evaluate the therapeutic effectiveness of antibodies in hamsters, Syrian golden hamsters were infected by intraperitoneal inoculation of 1000× LD_50_ NiV-M (corresponding to 8.55×10³ TCID_50_) per animal after being anesthetized with isoflurane. At 24 hours post-infection, the hamsters were intraperitoneally injected with antibody at a dosage of 10 mg per kilogram. In the prophylactic experiment, Syrian golden hamsters were administered a 10 mg kg^-1^ dose of antibody by the intraperitoneal route 24 hours before being intraperitoneally challenged with 1000× LD_50_ NiV-M (corresponding to 8.55×10³ TCID_50_). The control group was given an equivalent dose of isotype control antibody. All animals were observed for disease symptoms, weighed daily for 14 days after the challenge, and then observed for survival daily for another 14 days.

### Statistical Analyses

Statistical analyses were performed using GraphPad Prism software (v10). Details of statistical analysis can be found in figure legends and methods. Half maximal binding or neutralization concentrations were determined from serial dilution curves generated in GraphPad Prism; data is represented as mean with standard deviation (SD). Data from ADCC induction assays was collated from 3 independent experiments; mean and SD are shown. Statistical analysis of this data was conducted using a one-way ANOVA. Differences in survivability of animals was quantified via the log-rank Mantel-Cox test. Given that multiple comparisons were conducted, the α value of significance was adjusted using the Holm-Šídák correction method.

## Data Availability

### Lead contact

Further information and requests for resources and reagents should be directed to and will be fulfilled by the lead contact, Prof Daniel Watterson.

### Materials availability

Materials generated in this study are available upon request subject to a Materials Transfer Agreement.

### Data and code availability

Sequencing data generated from NiV immune escape experiments have been deposited to NCBI Sequence Read Archive and are available under accession PRJNA1022870. Generated cryoEM model has been deposited to Protein Data Bank (PDB 9B9E) and associated cryoEM maps have been deposited to Electron Microscopy Data Bank (EMD-44380, EMD-49520). This paper does not report original code. Any additional information required to reanalyse the data reported in this paper is available from the lead contact upon request.

