## Supplementary Figures for "A protective bispecific antibody targets both Nipah virus surface glycoproteins and limits viral escape"

Fig S1

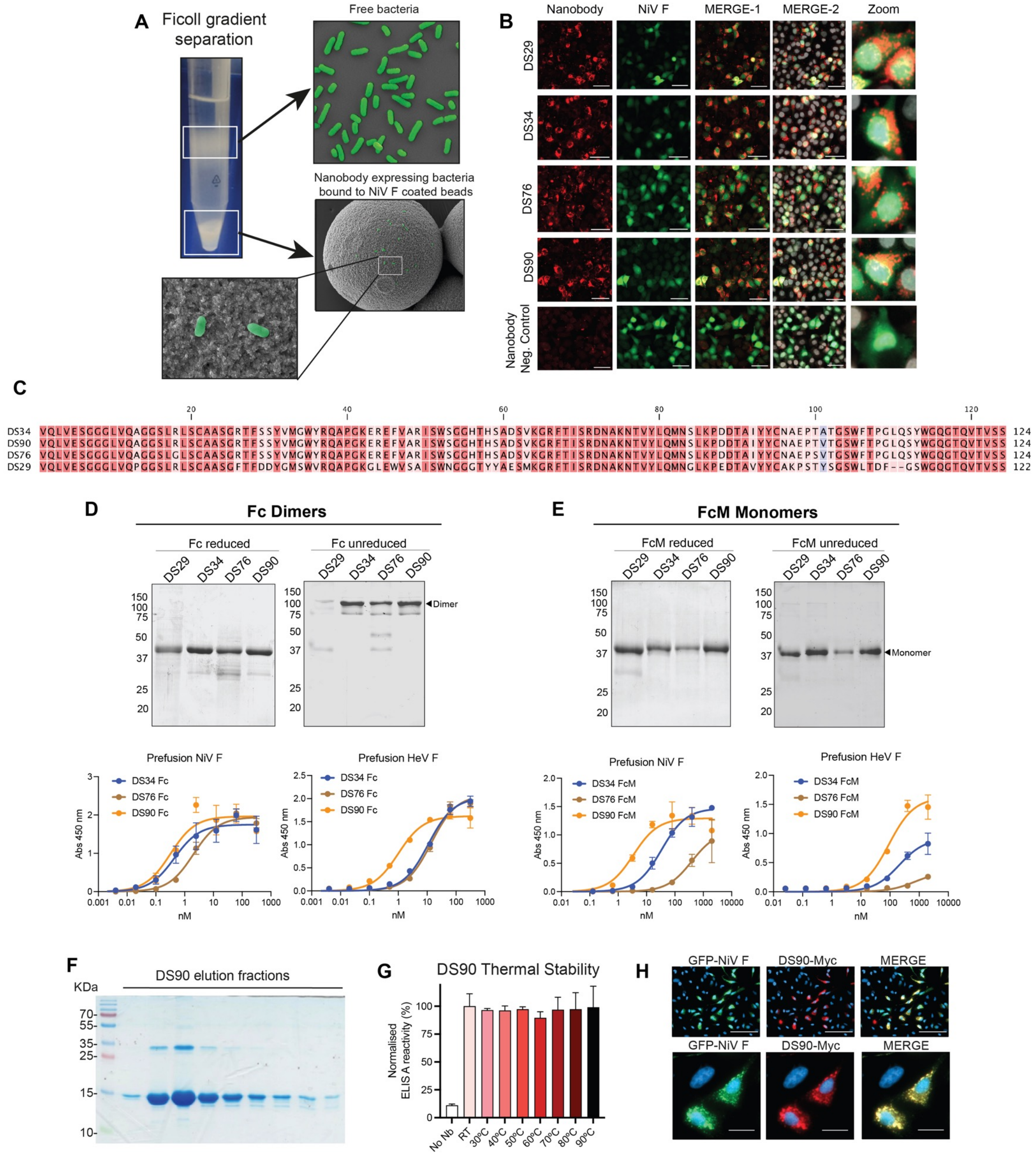
A
Ficoll gradient separation
Free bacteria
Nanobody expressing bacteria bound to NiV F coated beads
B
Nanobody
NiV F
MERGE-1
MERGE-2
Zoom
DS29
DS34
DS76
DS90
Nanobody Neg. Control
C
DS34 VQLVESGGGLVQAGGSLRLSCAASGRTFSSYVMGWYRQAPGKEREFVARISWSGGHTHSADSVKGRFTISRDNAKNTVYLQMNSLKPDDTAIYYCNAEPTATGSWFTPGLQSYWGQGTQVTVSS 124
DS90 VQLVESGGGLVQAGGSLRLSCAASGRTFSSYVMGWYRQAPGKEREFVARISWSGGHTHSADSVKGRFTISRDNAKNTVYLQMNSLKPDDTAIYYCNAEPTVTGSWFTPGLQSYWGQGTQVTVSS 124
DS76 VQLVESGGGLVQAGGSLGLSCAASGRTFSSYVMGWYRQAPGKEREFVARISWSGGHTHSADSVKGRFTISRDNAKNTVYLQMNSLKPDDTAIYYCNAEPSVTGSWFTPGLQSYWGQGTQVTVSS 124
DS29 VQLVESGGGLVQPGGSLRLSCAASGFTFDDYGMSWVRQAPGKGLEWVSAISWNGGGTYYAESMKGRFTISRDNAKNTVYLQMNGLKPEDTAVYYCAKPSTYSGSWLTDF--GSWGQGTQVTVSS 122
D
Fc Dimers
Fc reduced
Fc unreduced
DS29 DS34 DS76 DS90
Dimer
Prefusion NiV F
Prefusion HeV F
DS34 Fc
DS76 Fc
DS90 Fc
Abs 450 nm
nM
E
FcM Monomers
FcM reduced
FcM unreduced
Monomer
DS34 FcM
DS76 FcM
DS90 FcM
F
KDa
DS90 elution fractions
G
DS90 Thermal Stability
Normalised ELISA reactivity (%)
No Nb
RT
30°C
40°C
50°C
60°C
70°C
80°C
90°C
H
GFP-NiV F
DS90-Myc
MERGE

Fig S2

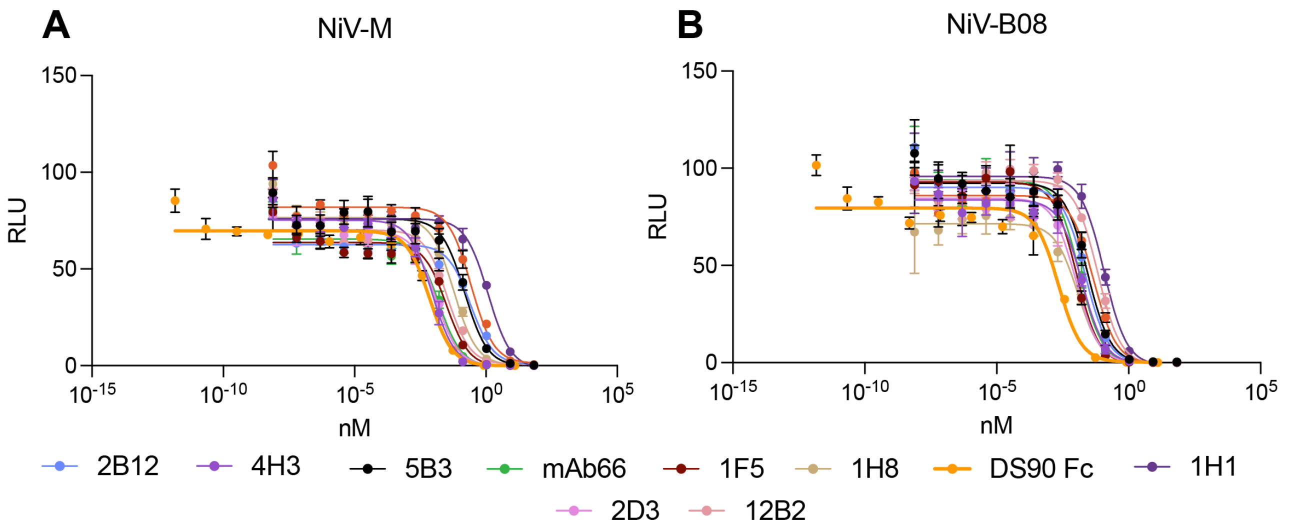


C

| | NiV-M | NiV-B08 |
|---|---|---|
| DS90 Fc | 0.007 | 0.002 |
| 4H3 | 0.009 | 0.017 |
| 1A9 | 0.271 | 0.047 |
| 1H1 | 1.203 | 0.116 |
| 2D3 | 0.012 | 0.010 |
| 2B12 | 0.260 | 0.023 |
| 12B2 | 0.034 | 0.067 |
| 1F5 | 0.031 | 0.010 |
| 5B3 | 0.154 | 0.028 |
| mAb66 | 0.016 | 0.013 |
| 1H8 | 0.060 | 0.012 |

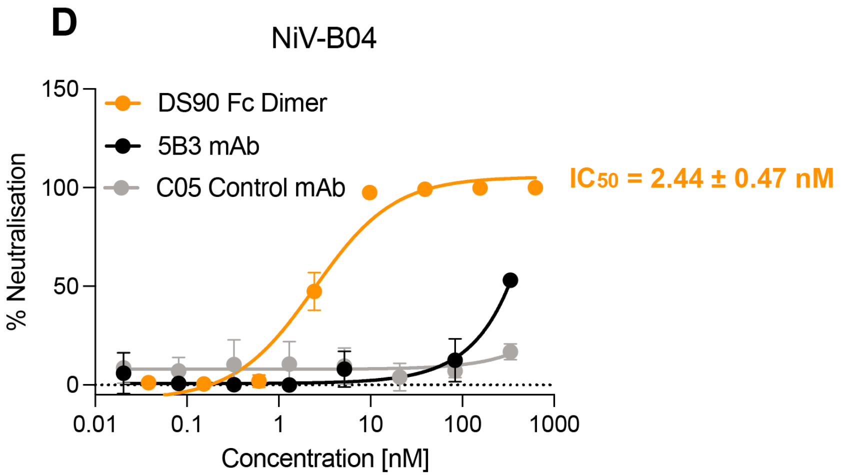

Fig S3

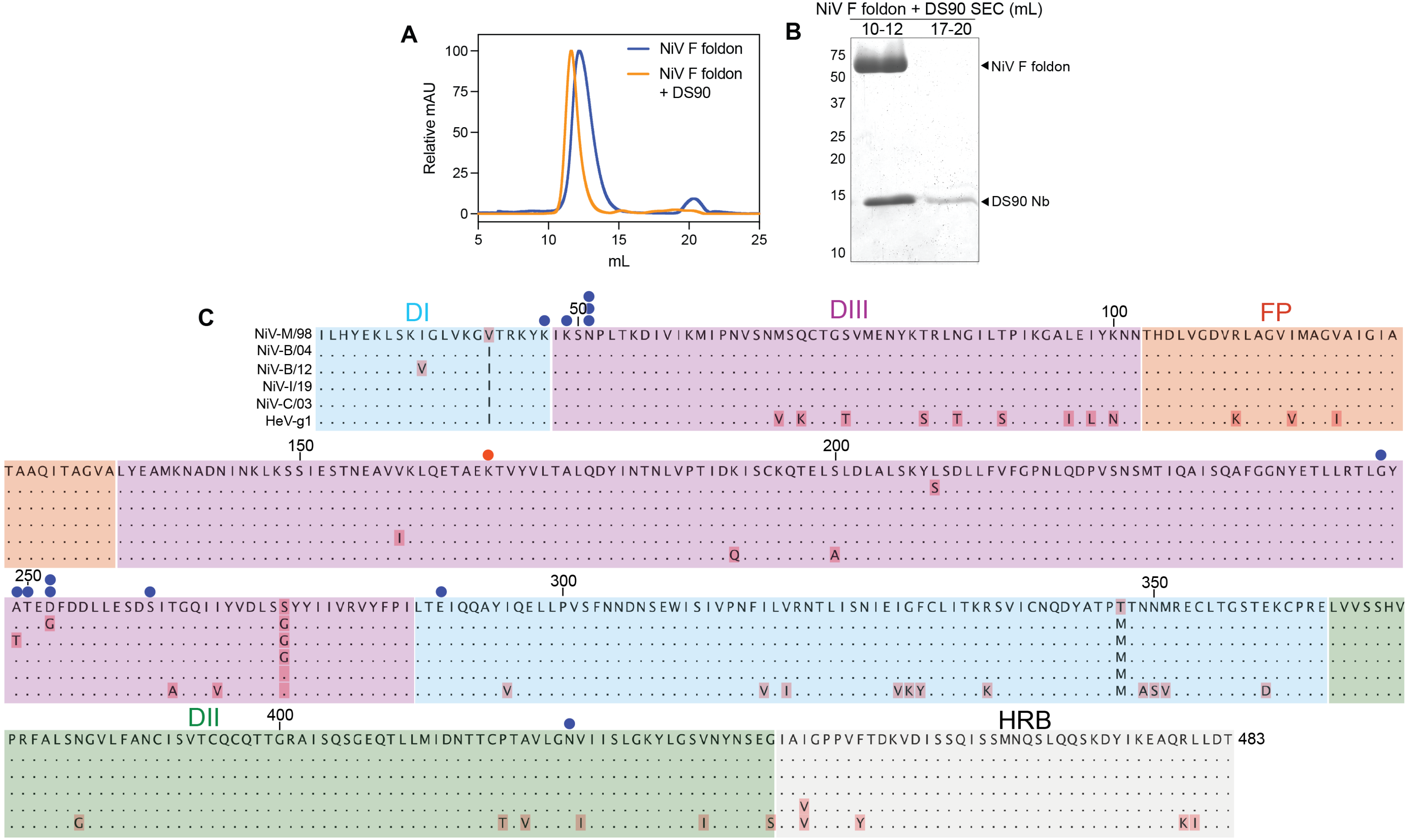

A
Relative mAU
mL
NiV F foldon
NiV F foldon + DS90
B
NiV F foldon + DS90 SEC (mL)
10-12
17-20
NiV F foldon
DS90 Nb
C
DI
DIII
FP
DII
HRB
NiV-M/98
NiV-B/04
NiV-B/12
NiV-I/19
NiV-C/03
HeV-g1
483

Fig S4

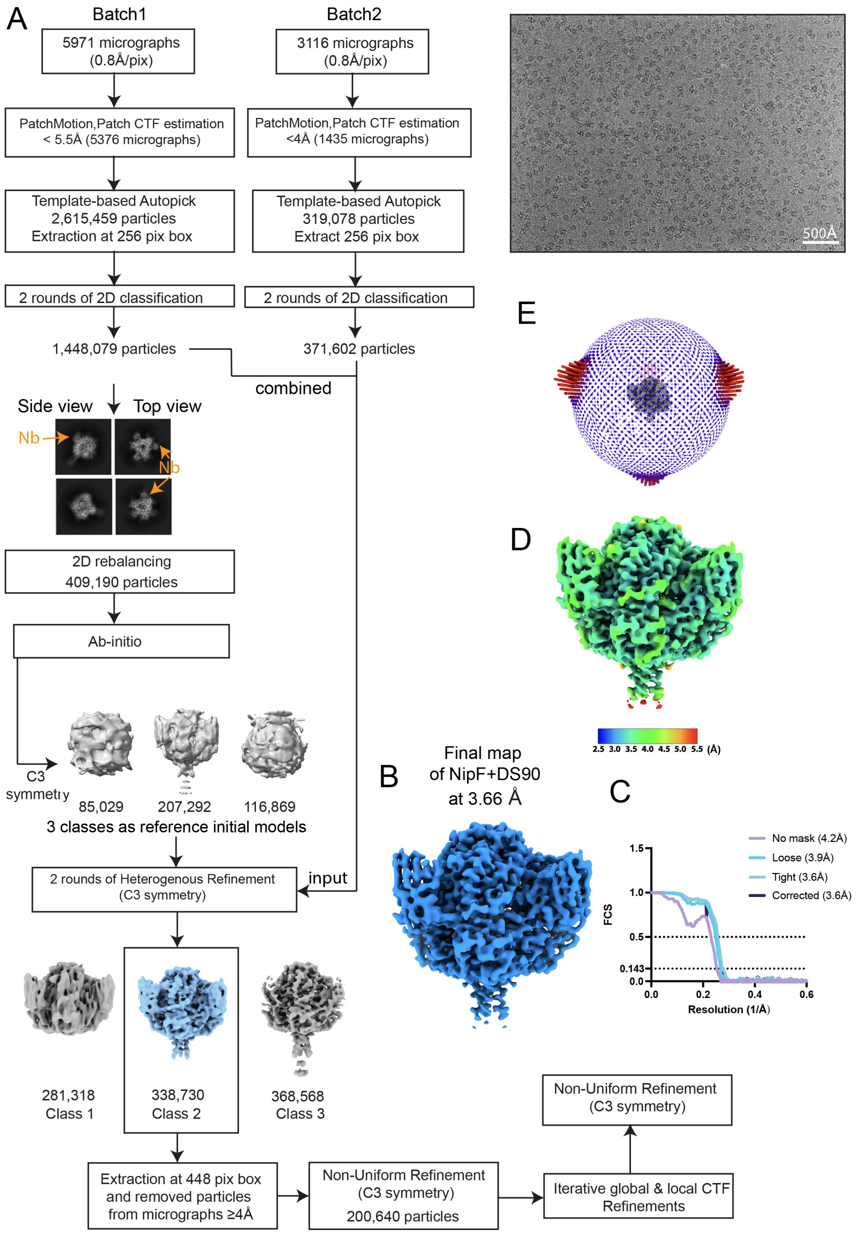

A
Batch1
5971 micrographs (0.8Å/pix)
PatchMotion,Patch CTF estimation < 5.5Å (5376 micrographs)
Template-based Autopick 2,615,459 particles Extraction at 256 pix box
2 rounds of 2D classification
1,448,079 particles
Batch2
3116 micrographs (0.8Å/pix)
PatchMotion,Patch CTF estimation <4Å (1435 micrographs)
Template-based Autopick 319,078 particles Extract 256 pix box
2 rounds of 2D classification
371,602 particles
combined
Side view
Top view
Nb
Nb
2D rebalancing 409,190 particles
Ab-initio
C3 symmetry
85,029
207,292
116,869
3 classes as reference initial models
2 rounds of Heterogenous Refinement (C3 symmetry)
input
281,318 Class 1
338,730 Class 2
368,568 Class 3
Extraction at 448 pix box and removed particles from micrographs ≥4Å
Non-Uniform Refinement (C3 symmetry) 200,640 particles
Iterative global & local CTF Refinements
Non-Uniform Refinement (C3 symmetry)
500Å
E
D
2.5 3.0 3.5 4.0 4.5 5.0 5.5 (Å)
B
Final map of NipF+DS90 at 3.66 Å
C
No mask (4.2Å)
Loose (3.9Å)
Tight (3.6Å)
Corrected (3.6Å)
FCS
1.5
1.0
0.5
0.143
0.0
0.0
0.2
0.4
0.6
Resolution (1/Å)

Fig S5

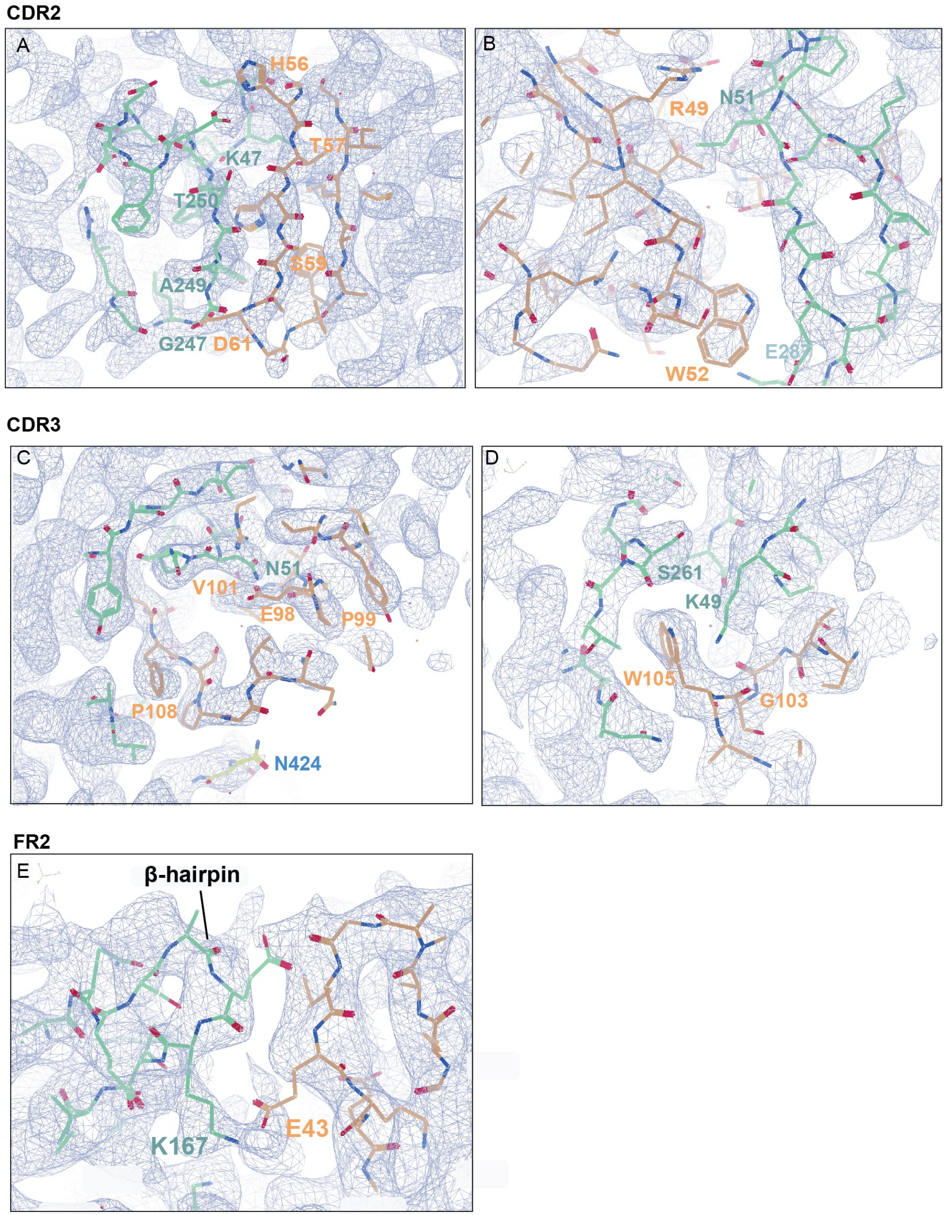
CDR2
A
H56
K47
T57
T250
S59
A249
G247
D61
B
R49
N51
W52
E287
CDR3
C
N51
V101
E98
P99
P108
N424
D
S261
K49
W105
G103
FR2
E
β-hairpin
K167
E43

#### Table S1

| | Nipah F protein in complex with DS90 (PDB ID: 9B9E , EMD-44380) |
|---|---|
| Data collection and processing | |
| Magnification | 50,000X |
| Voltage (kV) | 300 |
| Electron exposure (e– Å$^{-2}$) | 87.5 |
| Defocus range (μm) | −0.5 to −2.5 |
| Pixel size (Å) | 0.8 |
| Symmetry imposed | C3 |
| Initial particle images (no.) | 2,615,459 |
| Final particle images (no.) | 200,640 |
| Map resolution (Å) | 3.66 |
| FSC threshold | 0.143 |
| Map resolution range (Å) | 2.5-5.5 |
| Refinement | |
| Initial model used (PDB code) | 5EVM |
| Model resolution (Å) | 3.59 |
| FSC threshold | 0.143 |
| Map sharpening B factor (Å$^2$) | - |
| Model composition | |
| Non-hydrogen atoms | 13542 |
| Protein residues | 1740 |
| Ligands | 12 |
| R.m.s. deviations | |
| Bond lengths (Å) | 0.008 (0) |
| Bond angles (°) | 0.930 (2) |
| Validation | |
| MolProbity score | 1.05 |
| Clashscore | 1.14 |
| Poor rotamers (%) | 0 |
| Ramachandran plot | |
| Favored (%) | 96.53 |
| Allowed (%) | 3.47 |
| Disallowed (%) | 0 |
| Rama-Z (No. of residues analysed, Ramachandran plot Z-score, RMSD) | |
| Whole | N = 1728, -1.31 (0.18) |
| Helix | N= 480, -1.47 (0.21) |
| Sheet | N = 411, -0.14 (0.24) |
| Loop | N = 837, -0.74 (0.20) |

Table S2

| Donor | | Acceptor | | D-A Distance (Å) | Type |
|---|---|---|---|---|---|
| Nipah F | ASN 424 ND2 | DS90 | PRO 108 O | 3.1 | Hydrogen bond |
| | LYS 47 NZ | | HIS 56 NE2 | 2.9 | |
| | LYS 49 NZ | | GLY 103 O | 2.9 | |
| | ASN 51 N | | VAL 101 O | 3.4 | |
| | ASN 51 ND2 | | GLU 98 OE2 | 2.9 | |
| | ASN 51 ND2 | | PRO 99 O | 3.0 | |
| | THR 250 OG1 | | THR 57 O | 2.8 | |
| | ALA 249 N | | SER 59 O | 2.8 | |
| | LYS 167 NZ | | GLU 43 OE2 | 2.8 | Salt bridge |
| DS90 | ARG 49 NH2 | Nipah F | ASN 51 O | 2.9 | Hydrogen bond |
| | TRP 52 NE1 | | GLU 287 O | 2.9 | |
| | HIS 56 ND1 | | ASP 252 OD1 | 3.1 | |
| | HIS 56 ND1 | | ASP 252 OD2 | 2.7 | |
| | SER 59 N | | ALA 249 O | 3.0 | |
| | ASP 61 N | | GLY 247 O | 2.7 | |
| | VAL 101 N | | ASN 51 OD1 | 3.1 | |
| | TRP 105 NE1 | | SER 261 O | 3.0 | |

### Table S3

| Atom1 | | Atom2 | | VDW Overlap | Distance (Å) |
|---|---|---|---|---|---|
| DS90 | TRP 105 CD2 | Nipah F | LYS 49 HZ2 | 0.591 | 2.019 |
| | HIS 56 HD1 | | ASP 252 CG | 0.474 | 2.406 |
| | TRP 52 CH2 | | GLN 289 HE21 | 0.445 | 2.255 |
| | TRP 105 CD2 | | LYS 49 NZ | 0.337 | 2.898 |
| | HIS 58 NE2 | | TYR 248 HB3 | 0.296 | 2.344 |
| | ASP 61 H | | GLY 247 O | 0.293 | 1.727 |
| | TRP 52 CZ3 | | GLN 289 HE21 | 0.285 | 2.415 |
| | HIS 56 NE2 | | LYS 47 HZ1 | 0.256 | 1.984 |
| | TRP 105 CE2 | | LYS 49 NZ | 0.15 | 3.085 |
| | HIS 56 HD1 | | ASP 252 OD2 | 0.147 | 1.873 |
| | TRP 52 CH2 | | GLN 289 NE2 | 0.142 | 3.183 |
| | TRP 105 CD1 | | LYS 49 HZ2 | 0.141 | 2.559 |
| | TRP 105 CE2 | | LYS 49 HZ2 | 0.14 | 2.47 |
| | HIS 58 NE2 | | PHE 282 CE1 | 0.124 | 3.216 |
| | TRP 52 HD1 | | THR 286 OG1 | 0.123 | 2.377 |
| | ARG 49 HH21 | | ASN 51 HB3 | 0.118 | 1.882 |
| | PHE 106 CD2 | | ALA 133 HB3 | 0.094 | 2.606 |
| | ARG 49 HH21 | | ASN 51 CB | 0.046 | 2.654 |
| | GLU 98 OE2 | | ASN 51 HD21 | 0.043 | 1.977 |
| | LYS 42 CA | | GLU 166 OE1 | 0.023 | 3.097 |
| | LYS 42 HA | | GLU 166 OE1 | 0.015 | 2.405 |
| | TRP 105 CE3 | | LYS 49 HZ2 | -0.023 | 2.723 |
| | HIS 56 NE2 | | LYS 47 NZ | -0.032 | 2.897 |
| | TRP 105 CD1 | | LYS 49 NZ | -0.049 | 3.374 |
| | TRP 105 CE3 | | LYS 49 NZ | -0.078 | 3.403 |
| | GLY 41 O | | GLU 166 OE1 | -0.088 | 2.928 |
| | HIS 58 HD2 | | PHE 282 HE1 | -0.1 | 2.1 |
| | VAL 101 H | | ASN 51 OD1 | -0.1 | 2.12 |
| | TRP 105 CG | | LYS 49 HZ2 | 0.5 | 2.11 |
| | GLU 98 CD | | ASN 51 HD21 | 0.368 | 2.512 |
| | HIS 56 ND1 | | ASP 252 CG | 0.278 | 3.227 |
| | THR 57 O | | THR 250 HG1 | 0.22 | 1.8 |
| | ARG 49 NH2 | | ASN 51 HB3 | 0.183 | 2.442 |
| | SER 59 O | | ALA 249 H | 0.139 | 1.881 |

| Atom1 | | Atom2 | | VDW Overlap | Distance (Å) |
|---|---|---|---|---|---|
| DS90 | HIS 58 HE1 | Nipah F | TYR 248 CD1 | 0.131 | 2.569 |
| | TRP 105 CG | | LYS 49 NZ | 0.127 | 3.108 |
| | GLU 98 CD | | ASN 51 ND2 | 0.11 | 3.395 |
| | TRP 52 HE1 | | GLU 287 O | 0.105 | 1.915 |
| | THR 100 HA | | ASN 51 OD1 | 0.103 | 2.317 |
| | GLU 43 OE2 | | LYS 167 HZ1 | 0.103 | 1.917 |
| | TRP 52 CD1 | | THR 286 OG1 | 0.093 | 3.107 |
| | TRP 52 CZ2 | | GLN 289 HE21 | 0.068 | 2.632 |
| | TRP 105 CD1 | | LYS 49 HE3 | 0.055 | 2.645 |
| | HIS 58 NE2 | | TYR 248 CB | 0.053 | 3.287 |
| | HIS 58 NE2 | | PHE 282 CD1 | 0.05 | 3.29 |
| | ARG 49 HH21 | | ASN 51 O | 0.048 | 1.972 |
| | PRO 99 O | | ASN 51 HD22 | 0.047 | 1.973 |
| | HIS 58 CD2 | | PHE 282 CE1 | 0.043 | 3.357 |
| | HIS 58 CE1 | | TYR 248 HB3 | 0.038 | 2.662 |
| | TRP 52 CZ2 | | GLN 289 CG | 0.016 | 3.384 |
| | GLY 41 C | | GLU 166 OE1 | 0.014 | 3.016 |
| | TRP 52 CZ3 | | GLN 289 NE2 | 0.012 | 3.313 |
| | HIS 56 CE1 | | LYS 47 HZ1 | -0.005 | 2.705 |
| | TRP 105 O | | PRO 52 CG | -0.023 | 3.143 |
| | ARG 49 NH2 | | ASN 51 CB | -0.039 | 3.364 |
| | PHE 106 CG | | ALA 133 HB3 | -0.051 | 2.661 |
| | HIS 58 HD2 | | THR 250 CG2 | -0.051 | 2.751 |
| | TRP 105 CD1 | | LYS 49 CE | -0.061 | 3.461 |
| | HIS 56 ND1 | | ASP 252 OD2 | -0.061 | 2.706 |
| | HIS 56 CE1 | | LYS 47 NZ | -0.065 | 3.39 |
| | TRP 105 HE1 | | SER 261 O | -0.065 | 2.085 |
| | GLU 43 CD | | LYS 167 HD3 | -0.069 | 2.949 |
| | ASP 61 N | | GLY 247 O | -0.08 | 2.725 |
| | TRP 105 CD1 | | THR 263 CG2 | -0.093 | 3.493 |
| | SER 59 H | | ALA 249 O | -0.096 | 2.116 |
| | HIS 58 CD2 | | PHE 282 HE1 | -0.097 | 2.797 |

Fig S6

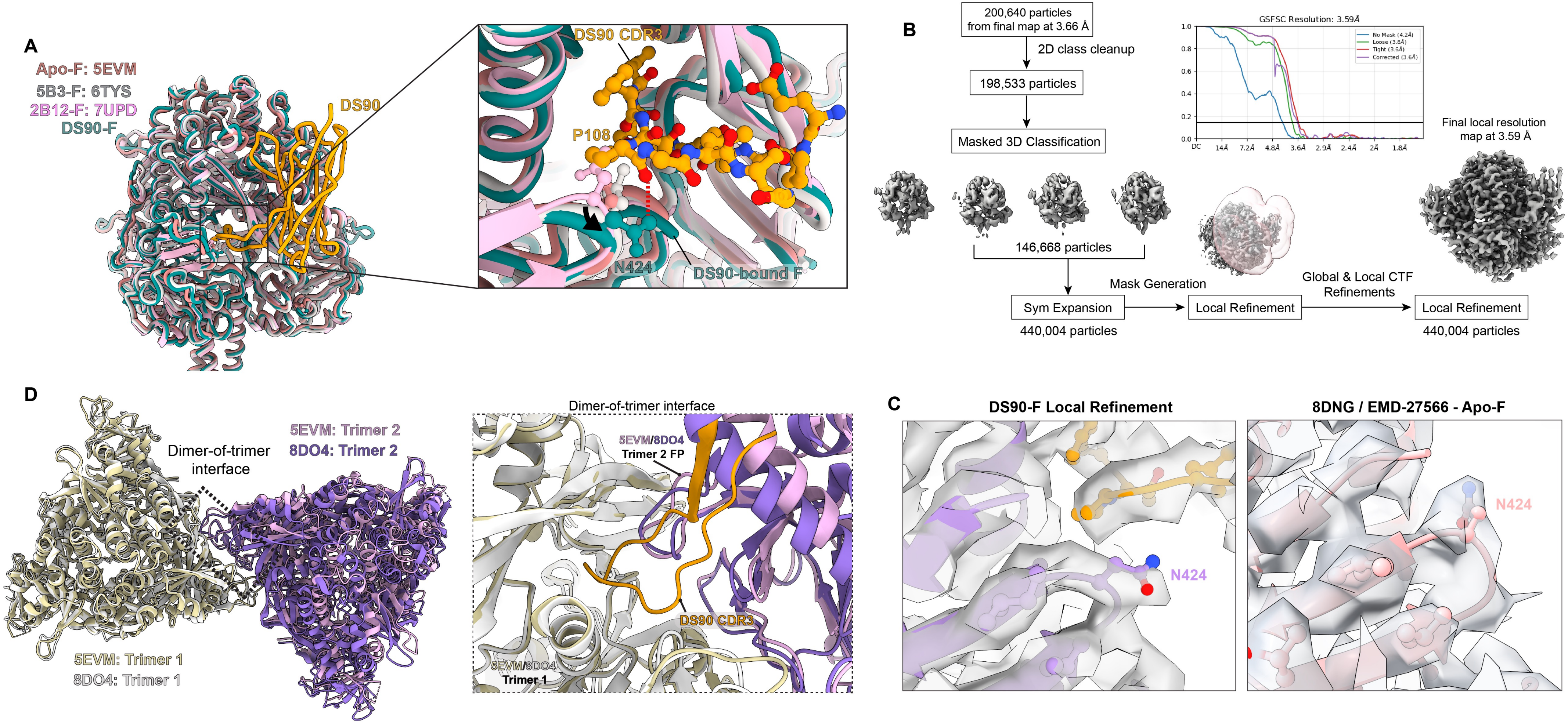

A
Apo-F: 5EVM
5B3-F: 6TYS
2B12-F: 7UPD
DS90-F
DS90
DS90 CDR3
P108
N424
DS90-bound F
B
200,640 particles
from final map at 3.66 Å
2D class cleanup
198,533 particles
Masked 3D Classification
GSFSC Resolution: 3.59Å
No Mask (4.2Å)
Loose (3.8Å)
Tight (3.6Å)
Corrected (3.6Å)
Final local resolution
map at 3.59 Å
146,668 particles
Mask Generation
Global & Local CTF
Refinements
Sym Expansion
440,004 particles
Local Refinement
Local Refinement
440,004 particles
D
Dimer-of-trimer
interface
5EVM: Trimer 2
8DO4: Trimer 2
5EVM: Trimer 1
8DO4: Trimer 1
Dimer-of-trimer interface
5EVM/8DO4
Trimer 2 FP
DS90 CDR3
5EVM/8DO4
Trimer 1
C
DS90-F Local Refinement
8DNG / EMD-27566 - Apo-F
N424
N424

Fig S7

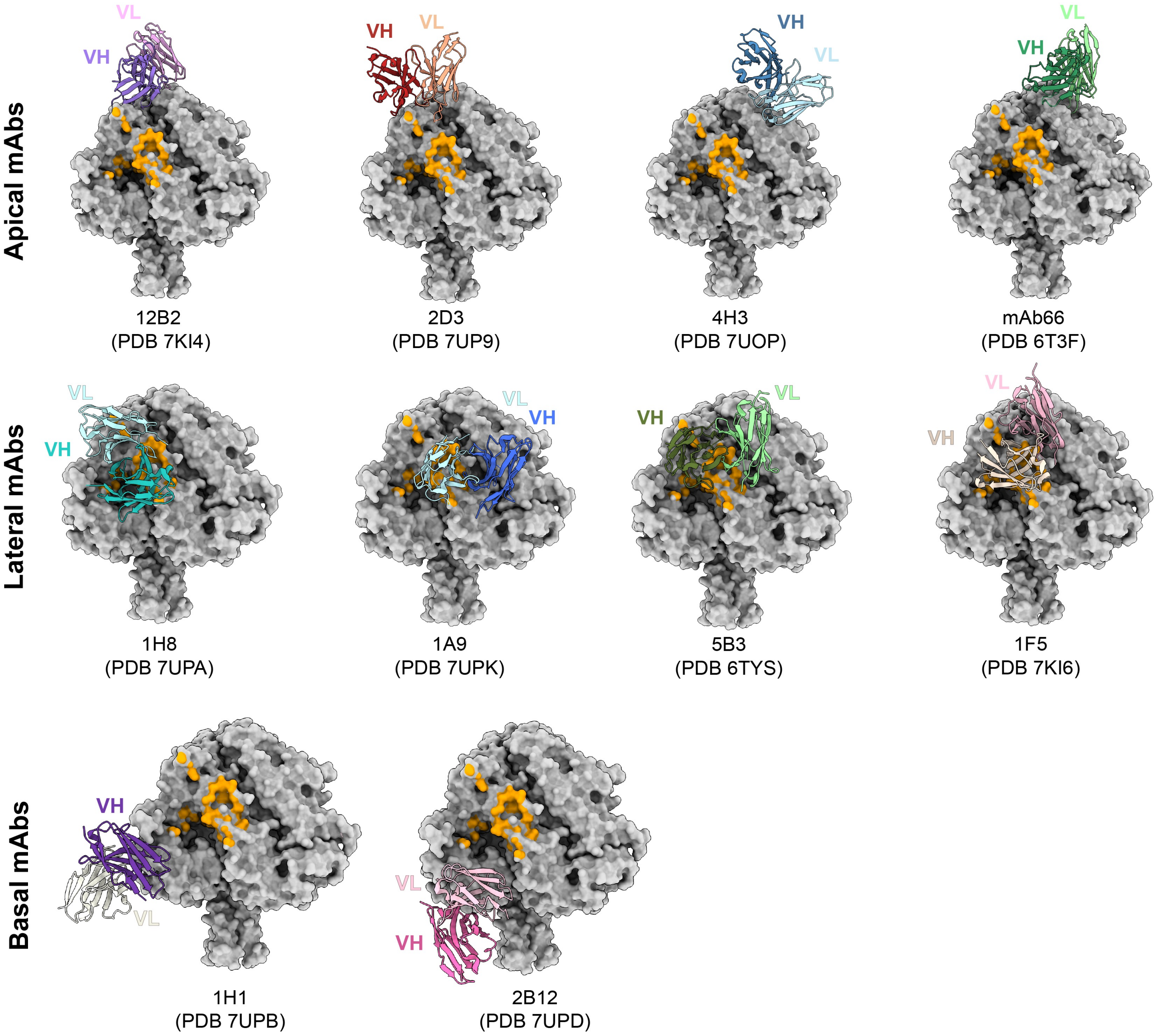

Apical mAbs
VL
VH
12B2
(PDB 7KI4)
VH
VL
2D3
(PDB 7UP9)
VH
VL
4H3
(PDB 7UOP)
VL
VH
mAb66
(PDB 6T3F)
Lateral mAbs
VL
VH
1H8
(PDB 7UPA)
VL
VH
1A9
(PDB 7UPK)
VH
VL
5B3
(PDB 6TYS)
VL
VH
1F5
(PDB 7KI6)
Basal mAbs
VH
VL
1H1
(PDB 7UPB)
VL
VH
2B12
(PDB 7UPD)

Fig S8

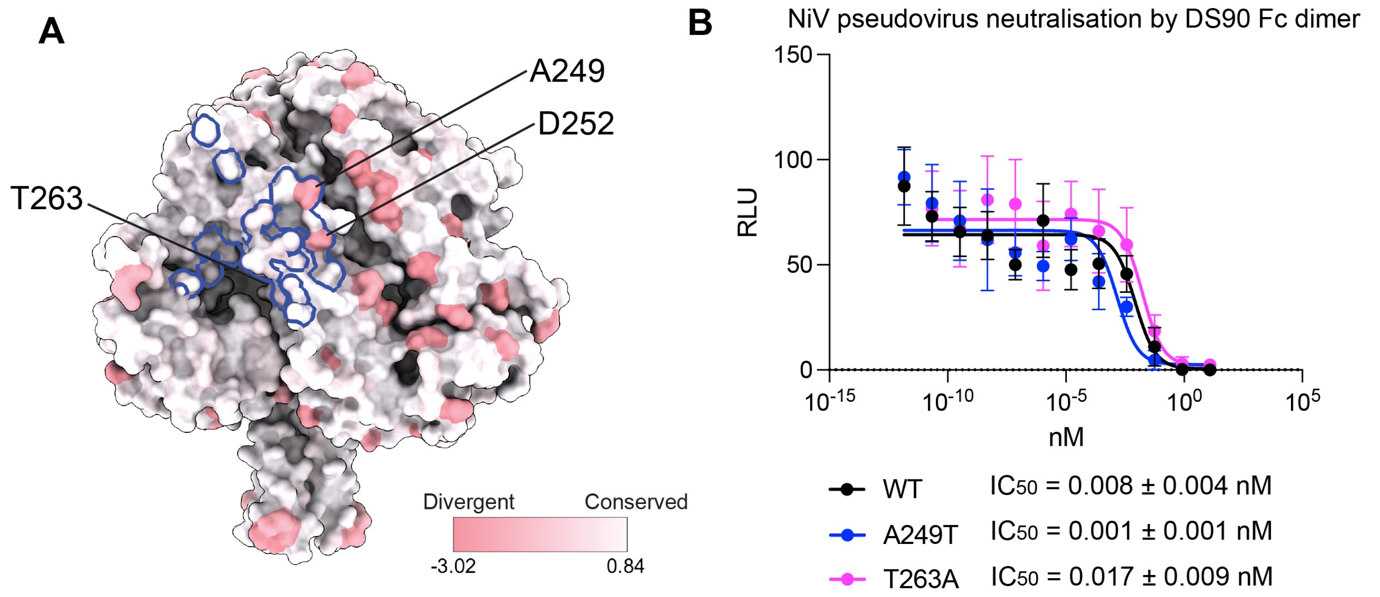

A
A249
D252
T263
Divergent
Conserved
-3.02
0.84
B
NiV pseudovirus neutralisation by DS90 Fc dimer
150
100
50
0
RLU
10-15
10-10
10-5
100
105
nM
WT
IC50 = 0.008 ± 0.004 nM
A249T
IC50 = 0.001 ± 0.001 nM
T263A
IC50 = 0.017 ± 0.009 nM

Fig S9

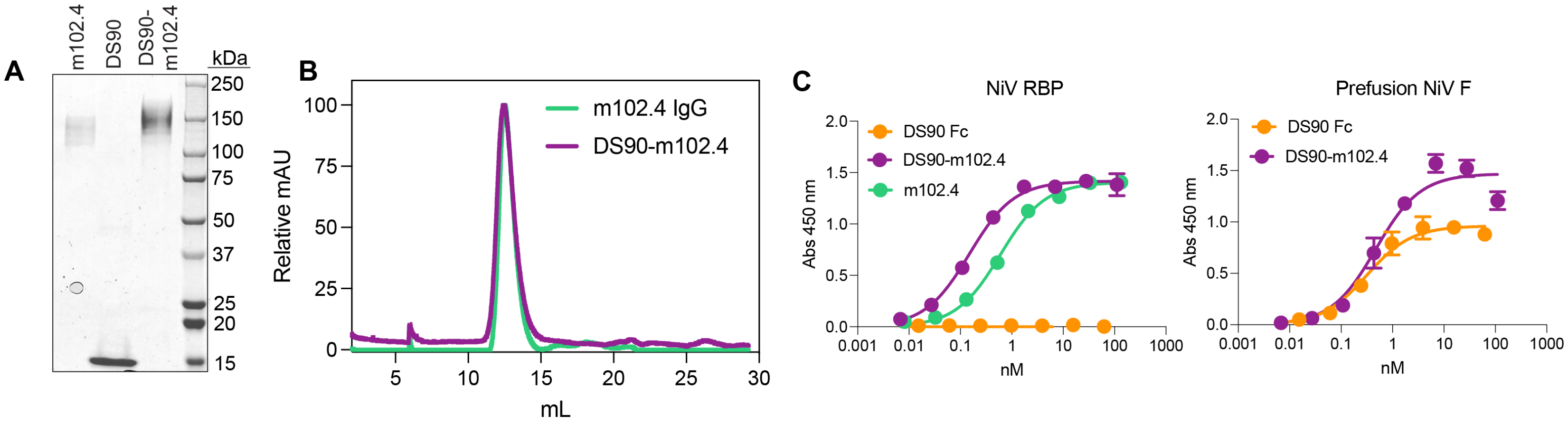


**D**

**Live Neutralisation CPE EPT (nM)**

| | DS90 Fc | DS90-m102.4 | m102.4 | Non-specific control |
|---|---|---|---|---|
| **NiV** | 5 | 2.2 | 83.33 | >2,500 |
| **HeV** | 9.8 | 8.6 | 333.3 | >2,500 |

Neutralisation

Fig S10

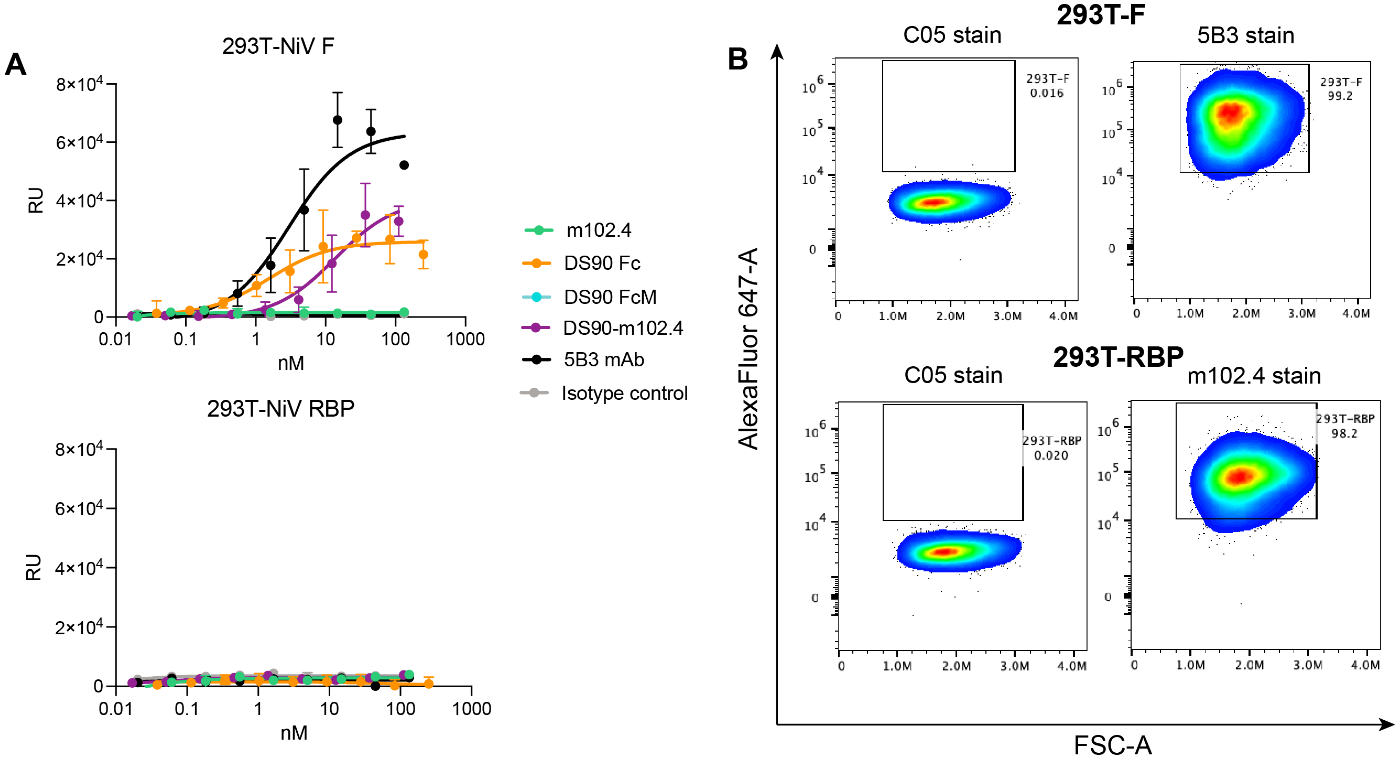
A
293T-NiV F
RU
nM
m102.4
DS90 Fc
DS90 FcM
DS90-m102.4
5B3 mAb
Isotype control
293T-NiV RBP
RU
nM
B
293T-F
C05 stain
5B3 stain
293T-F 0.016
293T-F 99.2
293T-RBP
C05 stain
m102.4 stain
293T-RBP 0.020
293T-RBP 98.2
AlexaFluor 647-A
FSC-A

#### Fig S11

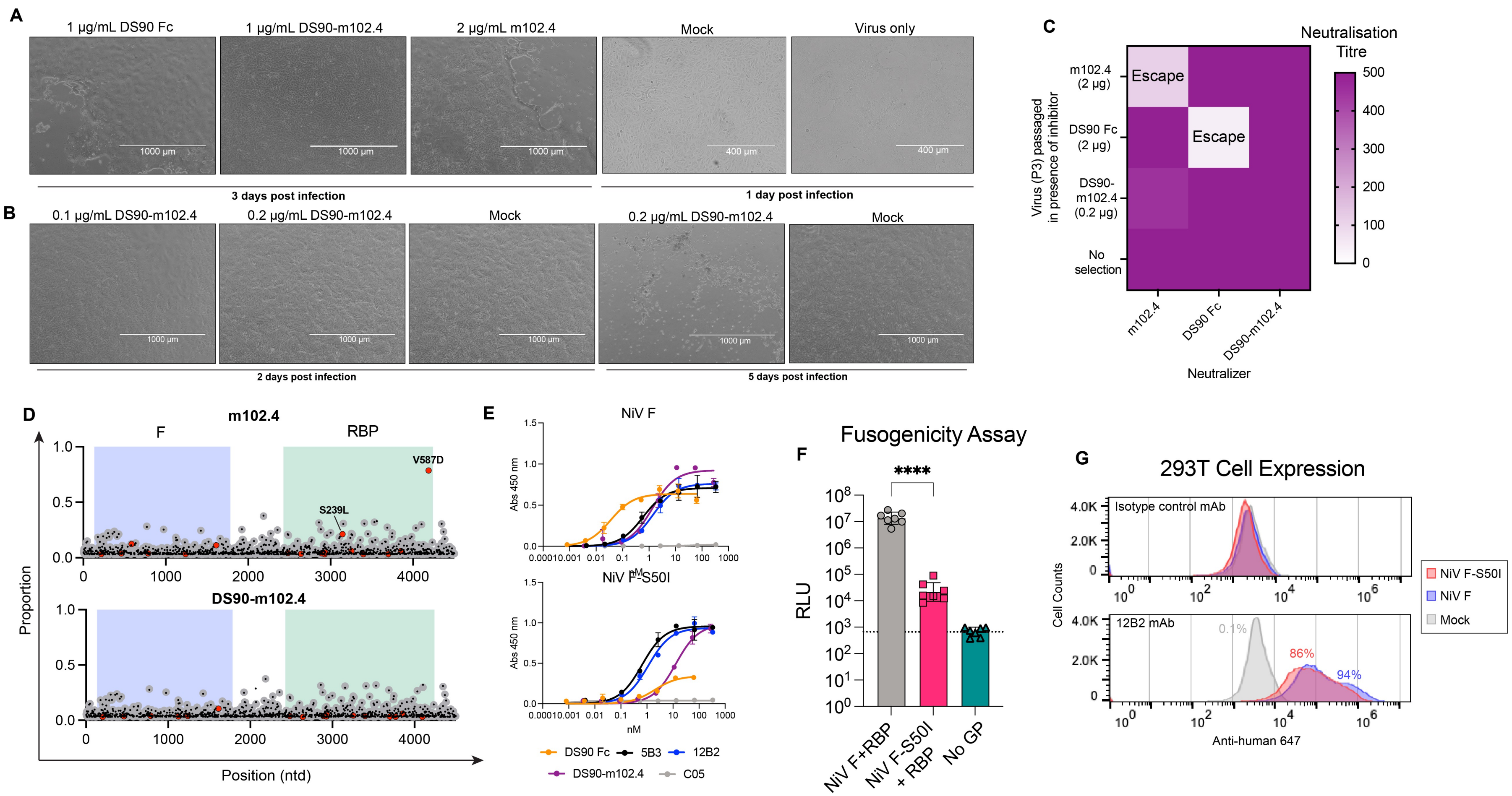
A
1 µg/mL DS90 Fc
1 µg/mL DS90-m102.4
2 µg/mL m102.4
Mock
Virus only
3 days post infection
1 day post infection
B
0.1 µg/mL DS90-m102.4
0.2 µg/mL DS90-m102.4
Mock
0.2 µg/mL DS90-m102.4
Mock
2 days post infection
5 days post infection
C
Neutralisation Titre
Virus (P3) passaged in presence of inhibitor
m102.4 (2 µg)
DS90 Fc (2 µg)
DS90-m102.4 (0.2 µg)
No selection
Escape
Escape
m102.4
DS90 Fc
DS90-m102.4
Neutralizer
D
m102.4
F
RBP
V587D
S239L
DS90-m102.4
Proportion
Position (ntd)
E
NiV F
NiV F-S50I
Abs 450 nm
nM
DS90 Fc
5B3
12B2
DS90-m102.4
C05
F
Fusogenicity Assay
RLU
****
NiV F+RBP
NiV F-S50I + RBP
No GP
G
293T Cell Expression
Isotype control mAb
12B2 mAb
Cell Counts
Anti-human 647
0.1%
86%
94%
NiV F-S50I
NiV F
Mock
